# SARS-CoV-2 spike-glycoprotein processing at S1/S2 and S2’and shedding of the ACE2 viral receptor: roles of Furin and TMPRSS2 and implications for viral infectivity and cell-to-cell fusion

**DOI:** 10.1101/2020.12.18.423106

**Authors:** Rachid Essalmani, Jaspreet Jain, Delia Susan-Resiga, Ursula Andréo, Alexandra Evagelidis, Rabeb Mouna Derbali, David N. Huynh, Frédéric Dallaire, Mélanie Laporte, Adrien Delpal, Priscila Sutto-Ortiz, Bruno Coutard, Claudine Mapa, Keith Wilcoxen, Etienne Decroly, Tram NQ Pham, Éric A. Cohen, Nabil G. Seidah

**Author notes:** Each of these authors contributed equally: Jaspreet Jain, Delia Susan-Resiga and Ursula Andreo.

## Abstract

The spîke (S)-protein of SARS-CoV-2 binds ACE2 and requires proteolytic “priming” at P**<u>R</u>**RA**<u>R</u>**_685_↓ into S1 and S2 (cleavage at S1/S2), and “fusion-activation” at a S2’ site for viral entry. *In vitro*, Furin cleaved peptides mimicking the S1/S2 cleavage site more efficiently than at the putative S2’, whereas TMPRSS2 inefficiently cleaved both sites. In HeLa cells Furin-like enzymes mainly cleaved at S1/S2 during intracellular protein trafficking, and S2’ processing by Furin at KPS**<u>KR</u>**_815_↓ was strongly enhanced by ACE2, but not for the optimized S2’ K<u>RR</u>KR_815_↓ mutant (μS2’), whereas individual/double KR815AA mutants were retained in the endoplasmic reticulum. Pharmacological Furin-inhibitors (Boston Pharmaceuticals, BOS-inhibitors) effectively blocked endogenous S-protein processing in HeLa cells. Furthermore, we show using pseudotyped viruses that while entry by a “pH-dependent” endocytosis pathway in HEK293 cells did not require Furin processing at S1/S2, a “pH-independent” viral entry in lung-derived Calu-3 cells was sensitive to inhibitors of Furin (BOS) and TMPRSS2 (Camostat). Consistently, these inhibitors potently reduce infectious viral titer and cytopathic effects, an outcome enhanced when both compounds were combined. Quantitative analyses of cell-to-cell fusion and spîke processing revealed the key importance of the Furin sites for syncytia formation. Our assays showed that TMPRSS2 enhances fusion and proteolysis at S2’ in the absence of cleavage at S1/S2, an effect that is linked to ACE2 shedding by TMPRSS2. Overall, our results indicate that Furin and TMPRSS2 play synergistic roles in generating fusion-competent S-protein, and in promoting viral entry, supporting the combination of Furin and TMPRSS2 inhibitors as potent antivirals against SARS-CoV-2.

**IMPORTANCE:** SARS-CoV-2 is the etiological agent of COVID-19 that resulted in >5 million deaths. The spike protein (S) of the virus directs infection of the lungs and other tissues by binding the angiotensin-converting enzyme 2 (ACE2) receptor. For effective infection, the S-protein is cleaved at two sites: S1/S2 and S2’. Cleavage at S1/S2, induces a conformational change favoring the recognition of ACE2. The S2’ cleavage is critical for cell-to-cell fusion and virus entry into host cells. Our study contributes to a better understanding of the dynamics of interaction between Furin and TMPRSS2 during SARS-CoV-2 entry and suggests that the combination of a non-toxic Furin inhibitor with a TMPRSS2 inhibitor could significantly reduce viral entry in lung cells, as evidenced by an average synergistic ∼95% reduction of viral infection. This represents a powerful novel antiviral approach to reduce viral spread in individuals infected by SARS-CoV-2 or future related coronaviruses.

## INTRODUCTION

Epidemics date from prehistoric times but are exacerbated by overcrowding and human impact on the ecosystem (1). The RNA coronaviruses (CoV) are zoonotic pathogens that occasionally spread in the human population, causing respiratory, enteric, renal, and neurological diseases (2). Electron microscopy of CoV revealed that the lipid envelope of each virion is surrounded by a “crown”-like structure (3), composed of multiple copies of a viral surface glycoprotein known as “spike” (S), which is essential for receptor binding and virus entry. Severe Acute Respiratory Syndrome coronavirus (SARS-CoV-1) and Middle East Respiratory Syndrome coronavirus (MERS-CoV) are infectious pathogenic viruses that appeared in humans at the beginning of the 21^st^ century (2, 4). At the end of 2019, a third CoV, namely SARS-CoV-2, emerged causing widespread respiratory and vascular illnesses (5), coined COVID-19 (6).

Like envelope glycoproteins of many infectious viruses (7–9), the secretory type-I membrane-bound S of SARS-CoV-2 is synthesized as a precursor (proS) that undergoes post-transcriptional cleavages by host cell proteases at specific sites to allow viral entry. During infection, the trimeric proS (monomer, 1,272 residues) is first processed at an S1/S2 cleavage site (Fig. 1A). Unlike SARS-CoV-1, the S-protein of SARS-CoV-2 exhibits an insertion of four critical amino acids (PRRA) at the S1/S2 junction (10–12), forming a canonical P**<u>R</u>**RA**<u>R</u>**_685_↓ Furin-like cleavage site (FCS). Such “priming” step divides the protein into two subunits S1 and S2 held together by non-covalent interactions. Following S-protein priming, the N-terminal S1-ectodomain undergoes a conformational change that exposes its receptor-binding-domain (RBD) (13), which recognizes the ACE2 entry receptor (11). The S2-subunit, which is responsible for the fusogenic activity of the spike-S glycoprotein, contains an additional “fusion-activation” proteolytic site (S2’) followed by an α-helical fusion peptide (FP) and two heptad-repeat domains (HR1 and HR2) preceding the transmembrane domain (TM) and cytosolic tail (CT) (Fig. 1A). It is thought that cleavage at S2’ triggers large-scale rearrangements, including a refolding step that is associated with the separation of S1- and S2-subunits and exposure of the hydrophobic α-helix FP, favoring fusion of viral and host cell membranes leading to virus entry (14). Fusion with host cells can occur either at the cell surface (pH-independent) or with internal membranes following endocytosis (pH-dependent) (15). However, the cognate host-cell proteases responsible for the S1/S2 and S2’ cleavages vary between coronaviruses and cell types (11, 12, 16-19).

**Figure 1:**
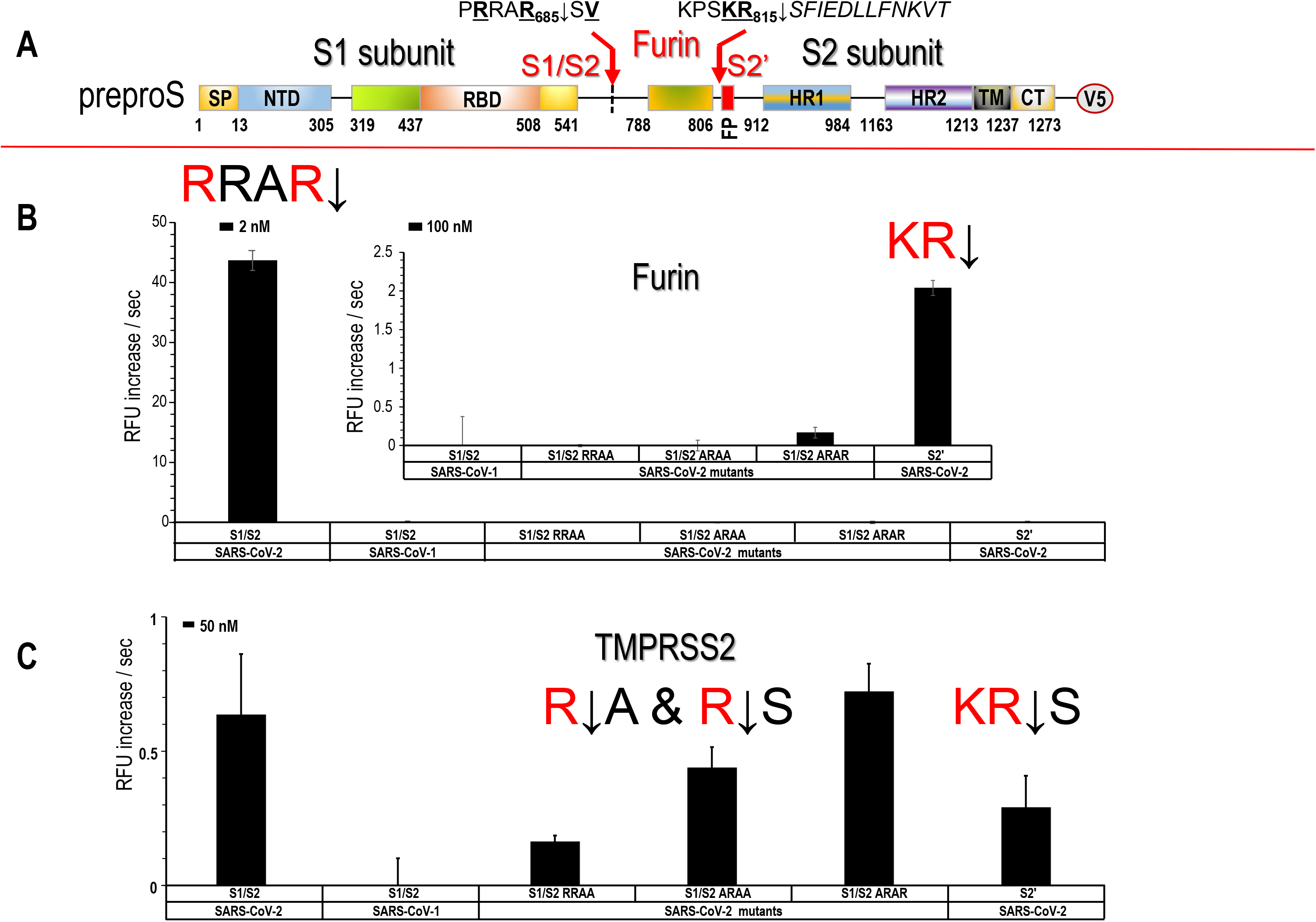
Processing of S-peptides by Furin and TMPRSS2. **(A)** Schematic representation of the primary structure of preproS and its domains and the predicted Furin-like S1/S2 site generating the S1- and S2-subunits, as well as the S2’ site preceding the fusion peptide (FP). The signal peptide (SP), N-terminal domain (NTD), receptor binding domain (RBD) to ACE2, the two heptad repeats HR1 and HR2, the transmembrane domain (TM), the cytosolic tail (CT) and the C-terminal V5-tag are indicated. **(B)** *In vitro* Furin activity against peptides mimicking the S1/S2 (and its mutants) and S2’ cleavage site sequence of the spike protein from SARS-CoV-2 and SARS-CoV-1, as described in SI-Table 1. Each substrate was tested at a final protease concentration of 2 and 100 nM. **(C)** *In vitro* TMPRSS2 activity (at 50 nM) against peptides mimicking the S1/S2 and S2’ cleavage site sequence of the spike protein from SARS-CoV-2 (and its mutants) described in SI-Table 1.

The proprotein convertases (PCs; genes *PCSKs*) constitute a family of nine secretory serine proteases that regulate various processes in both health and disease states (20). Through proteolysis, PCs are responsible for the activation and/or inactivation of many secretory precursor proteins, including virus/pathogen surface glycoproteins (9, 20). Seven PCs, including the widely expressed Furin, PC5A, PACE4 and PC7, cleave secretory substrates at specific single/paired basic amino acids (aa) within the motif (K/R)-X_n_-(K/R)↓, where Xn= 0, 2, 4 or 6 spacer X residues (20). Because of their critical functions, PCs, especially Furin (21), are implicated in many viral infections by inducing specific cleavages of envelope glycoproteins, a condition that allows not only the fusion of the viral lipid envelope with host cell membranes (9, 20), but can also lead to cell-to-cell fusion (syncytia), especially for viruses that undergo pH-independent fusion (22, 23). As the S1/S2 cleavage of SARS-CoV-2 is thought to play a critical role for cellular receptor recognition and virus entry, the efficacy and extent of this activation step by host proteases might be a key determinant regulating cellular tropism, viral pathogenesis, and human-to-human transmission. In contrast to SARS-CoV-1, the proS of SARS-CoV-2 contains a structurally exposed P**<u>R</u>**RA**<u>R</u>**_685_↓S**<u>V</u>** motif (10, 11) (Fig. 1A), which corresponds to a canonical FCS (9, 10, 20). This Furin-like motif is presumably cleaved during *de novo* virus egress (23) for S-protein priming and may play a key role for the efficient spread of SARS-CoV-2 to various human tissues compared to the more limited tropism of other lineage B β-coronaviruses (10, 24). Furthermore, based on the predicted S2’ KPS**<u>KR</u>**_815_↓S**F** sequence of SARS-CoV-2, we proposed (10) that Furin-like enzymes could also cleave at this S2’ site (Fig. 1A). Indeed, various reports have since supported the implication of Furin in the S1/S2 priming of the S-protein in human cell culture models (12, 25, 26) and *in vivo* in mice, hamsters and ferrets (27, 28). In addition, it was also suggested that the cell surface type-II transmembrane serine protease 2 (TMPRSS2) can enhance fusion by cleavage at S2’, but that S1/S2 cleavage is mostly Furin-dependent (18). The ability of the Arg/Lys-specific TMPRSS2 (29, 30) to directly cleave at S2’ was suggested based on the viral entry blockade by the TMPRSS2 inhibitor Camostat (31–33), and through silencing of TMPRSS2 expression using a morpholino oligomer (18), but direct evidence of its involvement in such spike protein processing at S2’ is still lacking. Thus, it is likely that one or more proteases regulate SARS-CoV-2 entry into human airway epithelial cells (18, 24). Furthermore, since the tissue-expression of TMPRSS2 is restricted to a limited set of cell types compared to that of the ubiquitously expressed Furin, the activity of the latter may widen viral tropism (34).

Thus, the major goals of the present study were to precisely define the respective roles of Furin and TMPRSS2 in the fusion activation, and to test the consequences of their inhibition on SARS-CoV-2 infectivity and S-mediated cell-to-cell fusion. Herein, using a multi-disciplinary approach, we provide mechanistic evidence supporting a critical role of the proprotein convertase Furin in the processing of SARS-CoV-2 spike protein. Specifically, we map the exact S2’ processing site by proteomics and highlight by mutagenesis the functional importance of S1/S2 and S2’ regions in viral entry and cell-to-cell fusion. For the first time, we demonstrate that three novel cell-permeable small molecules inhibitors of proprotein convertases developed by Boston Pharmaceuticals, referred hereafter as BOS-inhibitors, can potently inhibit proS processing at S1/S2 and S2’ by endogenous Furin-like proteases leading to efficient inhibition of viral entry, viral replication, and cell-to-cell fusion. Finally, our work sheds a new light on the role of TMPRSS2 in promoting ACE2 shedding and enabling S2’ processing, thereby leading to enhanced cell-to-cell fusion.

## RESULTS

### Comparative analysis of cleavage of SARS-CoV-2 peptides mimicking S1//S2 and S2’ processing sites by Furin and TMPRSS2

Furin is thought to be important in the processing of SARS-CoV-2 spike-glycoprotein (S) at the S1/S2 site (10, 24) while TMPRSS2 has been proposed to have an important role in activating S at S2’ (18, 31–33) (Fig. 1A). Nevertheless, the relative contributions of Furin and TMPRSS2 towards cleavage of SARS-CoV-2 S glycoprotein at both sites remain poorly defined. Thus, the susceptibility of SARS-CoV-2 S glycoprotein to Furin-cleavage was first assessed *in vitro*. Incubation of quenched fluorogenic peptides encompassing S1/S2 and S2’ sites (Supporting Information SI-Table 1) demonstrated that the S1/S2 site of SARS-CoV-2 S was efficiently cleaved by 2 nM Furin at pH 7.5 (Fig. 1B), whereas the S1/S2 site of SARS-CoV-1, which lacks an FCS, was not cleaved (Fig. 1B). Furin less efficiently cleaved the SARS-CoV-2-mimic peptide at S2’, requiring 50-fold higher enzyme concentrations (100 nM) to detect cleavage (inset Fig. 1B). The high specificity of the SARS-CoV-2 for processing at Furin-like motifs was next confirmed by demonstrating that substitutions of basic residues at the S1/S2 cleavage site (RRA**<u>A</u>**_685_↓S, **<u>A</u>**RA**<u>A</u>**_685_↓S, **<u>A</u>**RAR_685_↓S) dramatically impaired S1/S2 cleavage (Fig. 1B). Altogether, these data demonstrate that *in vitro* Furin best cleaves at S1/S2 and less efficiently at S2’. In contrast, TMPRSS2 did not efficiently cleave the S1/S2 and S2’ peptides (Fig 1C). The cleavage at S1/S2 became detectable only when TMPRSS2 was present at high concentration (50 nM). However, different from Furin, under this condition TMPRSS2 cleavage of peptides mimicking the S1/S2 Ala-mutants RRA<u>A</u>, <u>A</u>RA<u>A</u> and <u>A</u>RAR and S2’ was also evident (Fig. 1C). Taken together, these data emphasize the critical importance of the P1 and P4 Arg for Furin-mediated cleavage at S1/S2 and suggest that the likely Arg-motif recognized by TMPRSS2 is either (Ala/Arg)-Arg↓Ala or Ala-Arg↓Ser with a preference for Ala at P2 over Arg, and Ala or Ser at P1’.

### Furin and Furin-like proteases can process proS at S1/S2 and S2’ sites

To examine the ability of Furin and Furin-like enzymes to process the precursor proS of SARS-CoV-2 *in cellulo*, we used a HeLa cells model, which endogenously express Furin but not TMPRSS2 or ACE2 (*not shown*), as reported earlier (35). Here, we found that endogenous enzymes efficiently processed a V5-tagged proS (Fig. 1A), likely at the S1/S2 junction to generate a ∼100 kDa S2-like fragment (Fig. 2A). Interestingly, when the proprotein convertases PC5A, Furin, PC7 or PACE4 were transiently overexpressed, cleavage at this site became more prominent. Of note, a partial knockdown of Furin decreased S2 levels by more than 60% (SI-Fig. 1A). Furthermore, it was only when Furin or PC5A were overexpressed that cleavage at a potentially S2’ site was noticeable, yielding a ∼75 kDa fragment (Fig. 2A). The remaining ∼200 kDa proS_im_ corresponded to an immature precursor form that had not exited the ER, as attested by its sensitivity to both endoglycosidase-F and endoglycosidase-H (SI-Fig. 1B) and insensitivity to Furin-like convertases, which are only active in the TGN and/or cell surface/endosomes (20, 36).

**Figure 2:**
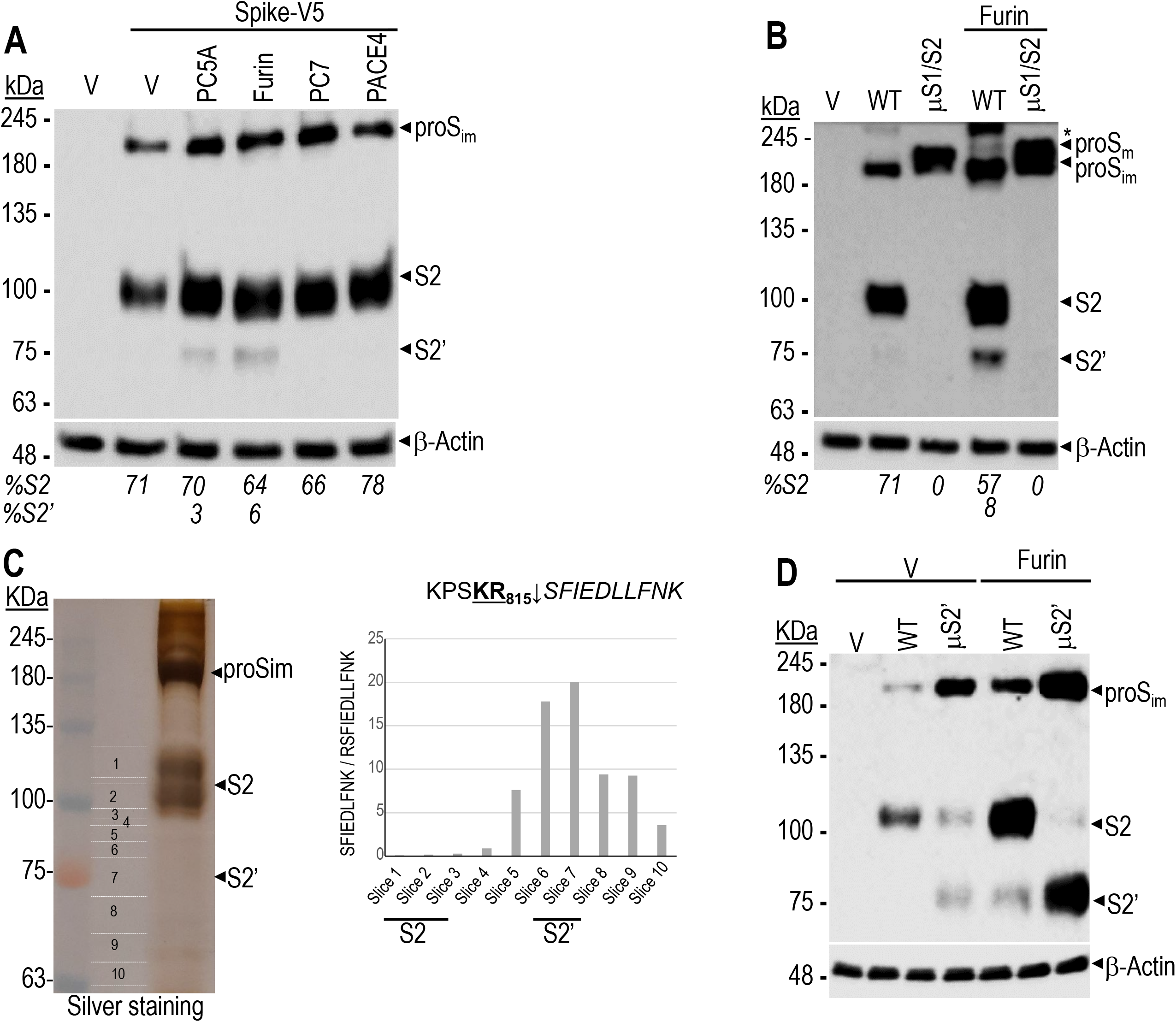
Processing of spike-glycoprotein in HeLa cells. (**A**) Western blot analyses of the processing of WT proS into V5-tagged S2 and S2’ by the proprotein convertases Furin, PC5A, PACE4 and PC7 following co-transfection of their cDNAs in HeLa cells. The migration positions of immature proS_im_, S2 and S2’ as well as the actin loading control are emphasized. V = empty pIRES-EGFP-V5 vector. (**B**) Western blot analyses of HeLa cells following co-transfection with cDNAs coding for either WT-S-protein or its double Ala-mutant [R685A + R682A] (μS1/S2) in the absence or presence of Furin cDNA at a ratio S:protease = 1:2. *Inconsistently observed oligomeric forms of proS. (**C**) Identification of S2’ cleavage site by MS/MS. WT-spike-glycoprotein was immunoprecipitated from HeLa cells using V5 agarose beads then resolved by SDS electrophoresis SDS/PAGE and subjected to silver staining (left panel); the positions of the slices are indicated (1 to 10). The MS/MS analysis of peptides generated by a Lys-specific protease (K_814_↓) are indicated; the data represent the ratio of SFIEDLLFNK825 to <u>R</u>_815_SFIEDLLFN<u>K</u>_825_ (right panel). (**D**) Western blot analyses of HeLa cells co-transfected with V5-tagged spike protein, WT (S) or its Furin-optimized S2’ (K<u>RR</u>KR_815_↓SF) mutant (µS2’), and empty vector (V) or Furin. (**A, B**) The estimated % cleavages into S1/S2 and S2’ are shown and were calculated as the ratio of the V5-immunoreactivity of the cleaved form to the sum of all forms. The data are representative of at least three independent experiments.

To precisely define the ∼100-kDa fragment, we mutated the S1/S2 site **<u>R</u>**RA**<u>R</u>**_685_↓S and found that the double Ala-mutant [**<u>A</u>**RA**<u>A</u>**_685_] (denoted μS1/S2) abrogated processing at S1/S2 and putative S2’ (Fig. 2B), highlighting once again the importance of the P4- and P1-Arg for recognition by Furin-like enzymes (20). The loss of Furin-like cleavage at S1/S2 resulted in accumulation of a higher molecular size band (∼230 kDa), representing mature proS, that exited the endoplasmic reticulum (ER), as confirmed by its resistance to endoglycosidase-H, while still sensitive to endoglycosidase-F digestion (SI-Fig. 1B).

To further define the Arg-residues critical for processing at S1/S2, we assessed the effect of single residue mutations: R682A, R685A and S686A and confirmed the critical importance of P1-Arg_685_ or P4-Arg_682_ for the generation of S2 by endogenous Furin (SI-Fig. 1C). However, unlike μS1/S2 (Fig. 2B), these single mutants were partially cleaved by overexpressed Furin (SI-Fig. 1C), reflecting the multi-basic nature of the S1/S2 recognition sequence and suggesting the importance of the P3 Arg_683_ (37). The S686A mutant was based on the prediction that Ser_686_ could be O-glycosylated (38), which may hamper processing at S1/S2 (39). However, like the WT-S, the S686A mutant was efficiently processed by Furin into S2 and S2’ (SI-Fig. 1C), suggesting the lack of O-glycosylation at Ser_686_ in HeLa cells.

We next used site-directed mutagenesis to identify the exact S2’ site cleaved by Furin. However, K814A, R815A and K814R815A mutants at P1 and P2 residues (20) of the predicted S2’ site (KPS<u>KR</u>_815_↓SF) altered S-protein trafficking resulting in a predominantly ER-retained proS_im_ protein, especially for the R815A and the double mutant (SI-Fig. 1D). Therefore, we resorted to mass spectrometry analysis of proteins migrating at the S2’ position to unambiguously identify this site. The peptides generated by a Lys-specific protease (K_814_↓) allowed for the discrimination between SFIEDLLFN<u>K</u>_825_ that would be generated if Furin cleaved at Arg_815_, and <u>R</u>_815_SFIEDLLFN<u>K</u>_825_ that would be derived from N-terminally extended proteins, e.g., S2. Proteomic data (Fig. 2C) revealed a >50-fold higher ratio of <u>S</u>FIEDLLFN<u>K</u>_825_ to <u>R</u>SFIEDLLFN<u>K</u>_825_ for the S2’ product, demonstrating that the N-terminus of S2’ starts at Ser_816_ and that Furin cleaves after Arg_815_↓ in the sequence KPSK**<u>R</u>**_815_↓SFIEDLLFNKVT (Fig. 1A). To further demonstrate the role of Furin in S2’ cleavage we generated a Furin-optimized S2’ site (called μS2’) with a polybasic sequence K<u>RR</u>KR_815_↓SF in proS and found that this derivative was very efficiently cleaved by endogenous and especially by overexpressed Furin, yielding a similar ∼75 kDa fragment (Fig. 2D). Altogether, these data demonstrate that the S2’ cleavage occurs at Arg_815_↓ and further reveal that this site can be partially processed by overexpressed Furin and/or PC5A.

### Processing at S2’ by Furin is enhanced in the presence of ACE2

Immunocytochemical analyses of HeLa cells co-expressing S or µS1/S2-S and ACE2 showed that both S-proteins and ACE2 co-localized at the cell surface independent of the state of proS processing (SI-Fig. 2). Given that binding of SARS-CoV-2 S-trimer to the dimeric ACE2 receptor has been proposed to trigger a conformational change in S1, promoting cleavage at S2’ (13, 40), we next examined whether this phenomenon would effectively increase/promote S2’ processing by Furin. To this end, we expressed the V5-tagged proS spike protein together with ACE2 and Furin in HeLa cells. We found that while not significantly affecting S1/S2 cleavage, ACE2 expression seemed to stabilize the S2 subunit and to strongly enhance the generation of S2’ by endogenous and overexpressed Furin (Fig. 3A). Amazingly, in the presence of ACE2, the μS1/S2-S, which is otherwise resistant to cleavage at S1/S2 by endogenous or overexpressed Furin (Fig. 2B), can be partially cleaved directly into S2’ by overexpressed Furin (Fig. 3B, last lane). Taken together, we conclude that binding of S-protein to ACE2 likely facilitates exposure of the S2’ site (41), thereby enhancing Furin processing at S2’.

**Figure 3:**
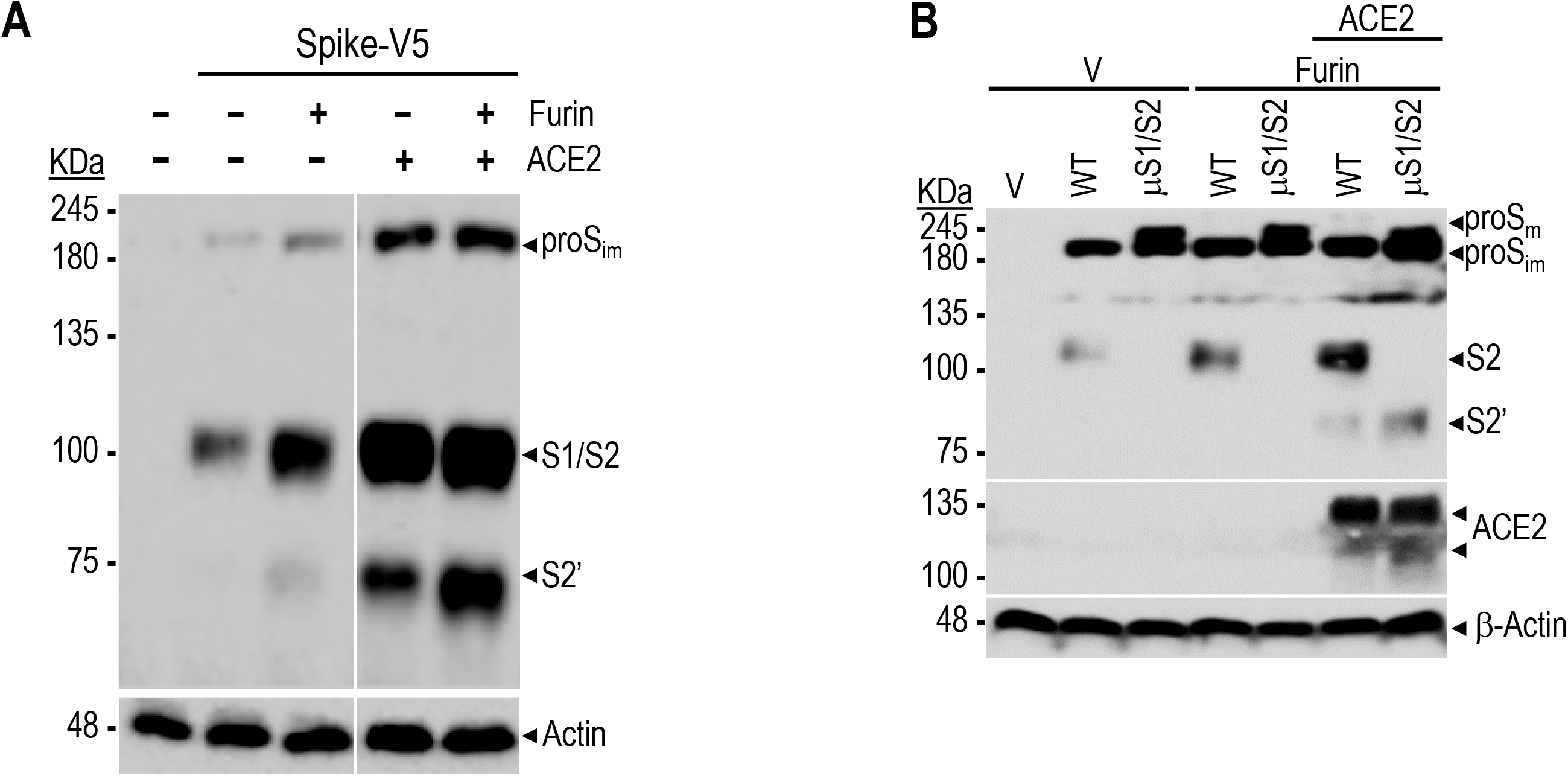
Processing of spike-glycoprotein at S2’ is enhanced in presence of ACE2. Western blot showing the impact of ACE2 on the processing of WT and μS1/S2 spike-glycoproteins by Furin. HeLa cells expressing empty vector (V), WT proS (**A**) or its μS1/S2 mutant (**B**) without or with Furin, ACE2 or both were analysed by Western blotting using anti-V5 antibody. The ratio of cDNAs used was S:ACE2:Furin = 1:1:1. The data are representative of at least three independent experiments.

### Furin-inhibitors block S1/S2 and S2’ cleavages

Given the importance of Furin in proS processing at the S1/S2 and S2’ sites, we next evaluated the activity of three novel non-toxic, cell-permeable Furin-like inhibitors developed by Boston Pharmaceuticals available as oral (BOS-981, BOS-318) or inhalable (BOS-857) formulations (Fig. 4A). Accordingly, we first tested *in vitro* the efficacy and selectivity of these inhibitors on purified soluble forms of Furin, PC5A, PACE4 and PC7 using a quenched fluorogenic substrate FAM-Q**<u>R</u>**V**<u>RR</u>**AVGIDK-TAMRA. As shown, the inhibitors effectively blocked substrate processing by all convertases with an IC_50_ of ∼7-9 nM compared to ∼9-10 nM for the known cell-permeable PC-inhibitor decanoyl-RVKR-chloromethylketone (dec-RVKR-cmk) (42, 43) (Fig. 4B). The Furin S1/S2 cleavage was also validated using a 12-residue quenched fluorogenic substrate DABSYL/Glu-TNSP**<u>RR</u>**A**<u>R</u>**↓SVAS-EDANS mimicking the S1/S2 priming site. The inhibition deduced after hill-plot curve fitting (Fig. 4C) gave an estimated IC_50_ of 4 ± 0.7 nM for BOS-981, 32 ± 4 nM for BOS-857 and 35 ± 5 nM for BOS-318. As well, BOS-inhibitors inhibited endogenous Furin-like processing of a dibasic bone morphogenic protein 10 (BMP10)-mimic (43) with an IC_50_ of ∼8 nM *versus* 5 nM for the dec-RVKR-cmk as determined by a cell-based Golgi imaging assay with U2OS cells (Fig. 4D). We further showed that BOS-inhibitors efficiently blocked S1/S2 and S2’ processing by endogenous Furin-like enzymes, resulting in a near complete inhibition at 0.3 μM, also obtained with 50 μM of dec-RVKR-cmk (RVKR; Fig. 4E). Overall, our data clearly demonstrate a role of Furin in the processing of proS at the S1/S2 and S2’ sites.

**Figure 4:**
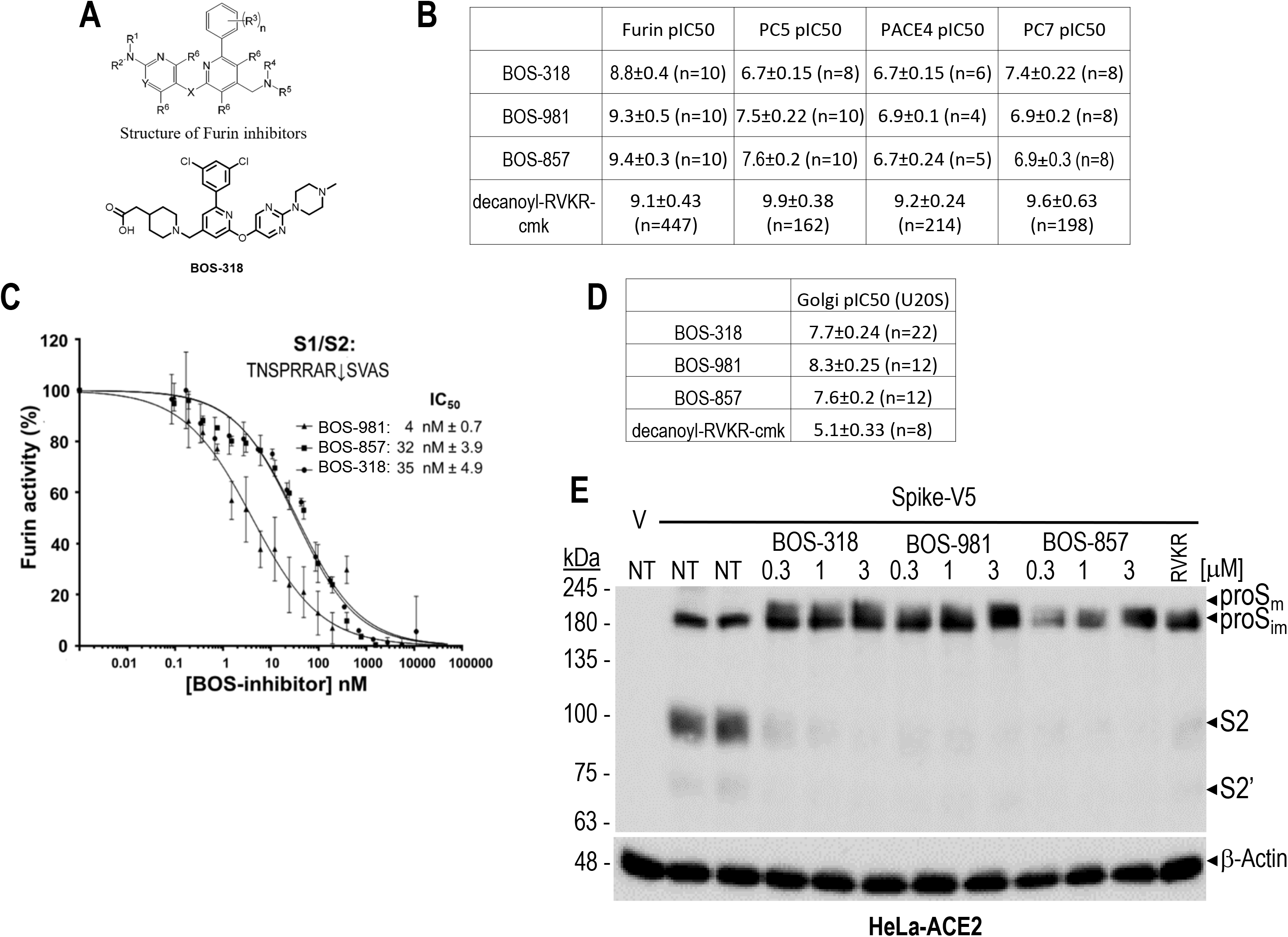
Inhibition of PCs by BOS compounds. (**A**) Chemical motif of BOS-inhibitors and representative structure of BOS-318. (**B**) *In vitro* BOS-inhibition of the cleavage of the fluorogenic dibasic substrate FAM-QRVRRAVGIDK-TAMRA by each of the proprotein convertases Furin, PC5 (PCSK5), PACE4 (PCSK6) and PC7 (PCSK7). All experiments were performed in 10 different wells and the average pIC_50_ (in nM) was calculated. Shown for comparison is the inhibitory pIC_50_ of the Furin-like inhibitor RVKR-cmk performed >100 times. (**C**) *In vitro* inhibition of Furin by the BOS compounds. Furin (2 nM) was incubated with increasing concentration of BOS-inhibitors, and its enzymatic activity against the synthetic peptides DABSYL/Glu-TNSP**<u>RR</u>**A**<u>R</u>**↓SVAS-EDANS (5 µM) was measured at pH 7.5 (n=3). (**D**) Golgi assay: table representing the effects of BOS-inhibitors on U2OS cells expressing each of Furin, PC5A, PACE4 and PC7 simultaneously transduced with a BacMam-delivered construct containing a Golgi-targeting sequence followed by a 12-amino acid Furin/PCSK cleavage site from Bone Morphogenic Protein 10 (BMP10) and GFP at the C terminus (GalNAc-T2-GGGGS-DSTARIRRNAKG-GGGGS-GFP). Dibasic cleavage releases NAKG-GGGGS-GFP thereby reducing the Golgi-associated fluorescence estimated by imaging. (**E**) Furin-inhibitors (BOS) abrogate endogenous processing of the spike-glycoprotein. Hela cells were transiently transfected with a cDNA encoding an empty vector (V) or with one expressing the V5-tagged spike (S) glycoprotein (spike-V5). At 5h pre-transfection, cells were treated with vehicle DMSO (NT, duplicate) or with the Furin-inhibitors at indicated concentrations, or RVKR-cmk at 50 μM. At 24h post-transfection media were replaced with fresh ones lacking (NT) or containing the inhibitors for an additional 24h. Cell extracts were analyzed by Western blotting using a mAb-V5. All data are representative of at least three independent experiments.

### Furin-like inhibitors reduce virus production in SARS-CoV-2-infected cells

We next examined whether blocking the processing of proS by BOS-inhibitors modulates SARS-CoV-2 infection. Indeed, in lung derived Calu-3 cells pretreated with 1 μM BOS-inhibitors for 24h before infection, we observed significantly decreased viral titers at 12, 24 and 48h post-infection (Fig. 5A). Importantly, the inhibitory effect was dose-dependent, reducing viral burden up to >30-fold with 1 µM BOS-318 (Fig. 5B; left panel). As well, the IC_50_ and selectivity index (44) of BOS-318 were 0.2 µM and 475, respectively (Fig. 5B; right panel). Importantly, the levels of spike (full length and cleaved S) and nucleocapsid proteins in the supernatant and cells were decreased in a dose-dependent manner (Fig. 5C), underscoring the crucial role played by Furin-like convertases in SARS-CoV-2 infection in this lung epithelial cell model. A similar analysis with BOS-857 and BOS-981 revealed comparable antiviral effects and selectivity index (SI-Figs. 3A, B). In addition, BOS-inhibitors were also evaluated in Vero E6 cells where SARS-CoV-2 entry and infection is established primarily *via* the endocytic pathway (11, 24). In this system and as expected, treatment with BOS-inhibitors led to weaker effect since we observed a decreased virus production by only ∼2.6-5.7-fold (SI-Fig. 4), possibly reflecting a role of Furin-like activity in early endosomes (36) for pH-dependent virus entry in Vero cells.

**Figure 5:**
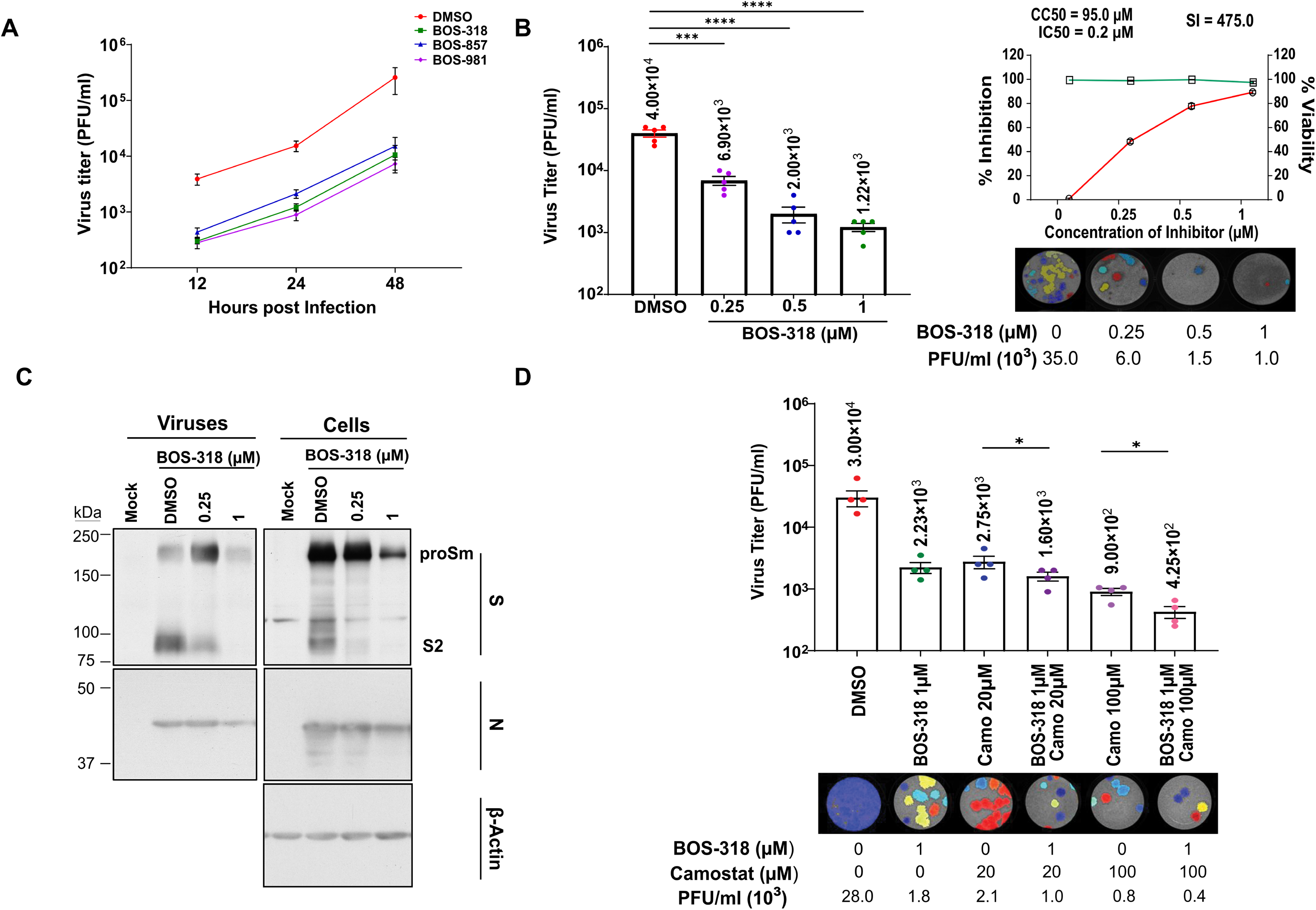
Furin-like inhibitors and Camostat treatment decrease SARS-CoV-2 infection in Calu-3 Cells. (**A**) Replication kinetics was studied at 12, 24 and 48h post-infection by plaque assay to determine PFUs of SARS-CoV-2 virus in the supernatant of infected Calu-3 cells treated or not with 1µM BOS-318, BOS-857 and BOS-981. A line graph represents results of the triplicate plaque assay results (mean ± SD). (**B**) The virus titers (PFU per milliliter) released in the supernatant (24h post-infection) of infected Calu-3 cells treated with indicated concentrations of BOS-318 were determined by plaque assay (mean ± SD of triplicates, *p < 0.05; **p < 0.01; ***p < 0.001) (left panel). The selectivity index (SI) of BOS-318 in Calu-3 cells as shown in top right panel was determined by CC_50_/IC_50_. The left y axis indicates the inhibition of virus titer (percent) relative to that of the untreated control group (red). The right y axis indicates the cell viability (percent) relative to that of the untreated control group (green). The CC_50_ (50% cytotoxic concentration), IC_50_ (half maximal inhibitory concentration), and SI (selectivity index) values for each inhibitor are as shown. Representative plaque images of infected Calu-3 cells treated with indicated doses of BOS-inhibitors are shown in the bottom right panel. (**C**) Immunoblots for the infected Calu-3 cells (right panel) and viral particles secreted in the supernatant (left panel) with and without treatment with BOS-inhibitors indicate reduced viral protein levels. Immunoblots were probed for the full-length (proSm) and cleaved (S2) fragments of viral S protein and nucleocapsid (N) protein as indicated; β-Actin was included as the loading control for the cells. (**D**) The virus titers (PFU per milliliter) released in the supernatant (24h post-infection) of infected Calu-3 cells treated with BOS-318 and/or Camostat (Camo) were determined by plaque assay (mean ± SD of duplicates, *, p < 0.05; **, p < 0.01; ***, p < 0.001) (top panel). Representative plaque images of infected Calu-3 cells are shown in the bottom panel. Color plaques differentiate the lawn (one color gray per well) from individual plaques (independent colors).

Since TMPRSS2 has been proposed to be important for viral entry at the plasma membrane, we next determined whether combining BOS-inhibitors and the TMPRSS2-inhibitor Camostat would have a synergistic effect, leading to a more pronounced antiviral effect in Calu-3. As shown, although these compounds could reduce viral replication individually, their co-treatment resulted in a synergistic inhibition of ∼95 ± 2.5% (>70-fold) of progeny infectious viruses (Fig. 5D and SI-Fig. 5), reinforcing the importance of both Furin-like proteases and TMPRSS2 in promoting efficient SARS-CoV-2 infection of Calu-3 cells.

### Furin-like inhibitors reduce viral entry by blocking processing of proS during biosynthesis and at the viral entry site

The more dramatic impact observed with Furin-like inhibitors on virus infection of Calu-3 cells *versus* Vero cells suggests that these inhibitors affect mainly the pH-independent entry mechanism. Thus, we next assessed the effect of BOS-inhibitors on viral entry. Using nanoluciferase-expressing HIV particles pseudotyped with WT, µS1/S2 or µS2’ S-proteins, we observed that the viral entry of µS1/S2-S pseudovirions is ∼10-fold reduced in Calu-3 cells (Fig. 6A). In contrast, all three pseudotyped viruses were at least 10-fold more infectious in HEK293T-ACE2 cells, suggesting that S-priming at S1/S2 is required for optimal viral entry in Calu-3 cells, but dispensable or perhaps even less desirable in HEK293T-ACE2 cells (Fig. 6A). Since these findings were similar to those in Vero E6 cells (45), we surmise that viral entry in HEK293T-ACE2 cells was through the same pH-dependent, endocytic route as reported for Vero cells (46, 47). This agrees with the fact that HEK293 cells allow endocytosis of SARS-CoV-2 pseudovirions *via* clathrin-coated vesicles (48). In the case of μS2’-S-expressing viral particles, entry was more efficient in both cell types (compare the same-coloured dots between WT-S and μS2’-S in absence of BOS, Fig. 6A), implying that S cleavage at the S2’optimized Furin-like site could enhance viral entry. When BOS-318 was present during biosynthesis of pseudovirions, processing of WT-S and μS2’-S was blocked (Fig. 6B), leading to reduced viral entry in Calu-3 by ∼3.6-to ∼12.5-fold, respectively (Fig. 6A). Thus, BOS-318 treatment phenocopied the effect of the µS1/S2 in both cell types. Nevertheless, under this condition viral entry in HEK293T-ACE2 cells was enhanced by ∼10-fold for WT-S and ∼2-fold for μS2’-S (Fig. 6A), suggesting that viral entry by the pH-dependent pathway does not require FCS processing.

**Figure 6:**
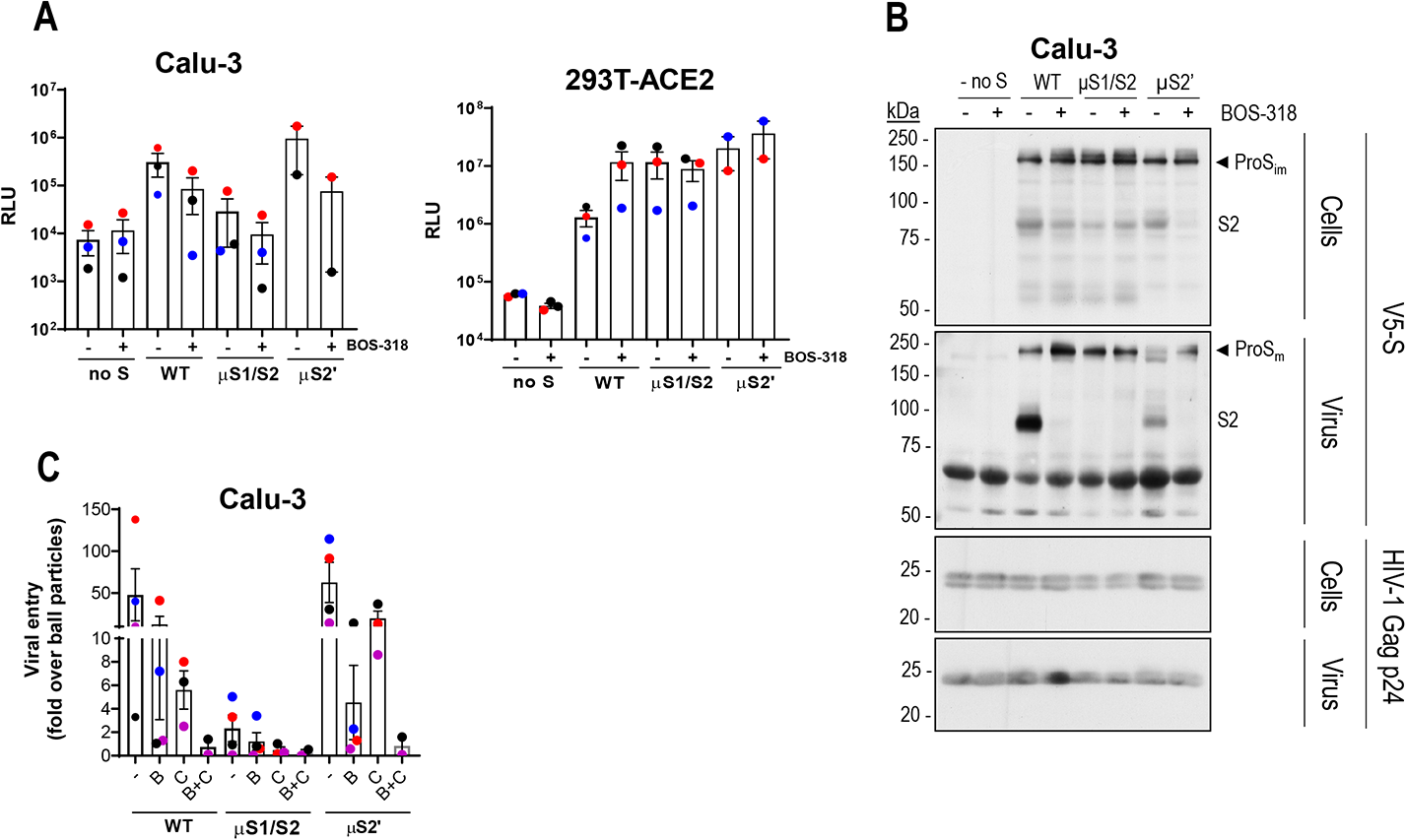
Processing of SARS-CoV-2 S by Furin-like convertases and TMPRSS2 is critical for viral entry in human lung epithelial cells but not in model HEK293 cells stably expressing ACE2. **(A)** Furin cleavage of proS at the S1/S2 site is required for SARS-CoV-2 pseudoviral entry in Calu-3 cells but not HEK293T-ACE2 cells. Cells were inoculated with nanoluciferase-expressing HIV particles pseudotyped with SARS-CoV-2: wild-type spike (WT), double Ala-mutant spike (µS1/S2) or Furin-optimized spike (μS2’). Inhibition of proS processing at S1/S2 by a novel Furin-like inhibitor (BOS-318) during pseudovirion packaging prevents viral entry in Calu-3 cells but not in HEK293T-ACE2 cells. (**B**) Western blot analyses show inhibition by BOS-318 of proS processing at S1/S2 site. Purified pseudovirions and cellular extracts of producing HEK293-T17 cells treated or not with BOS-318 inhibitor were separated on SDS-PAGE gel and analyzed for HIV-1 p24 and V5-tagged S-protein (proSm or cleaved, S2) as indicated. (**C**) Pre-treatment of Calu-3 cells with (B) 1 μM BOS-318, (C) 100 μM Camostat or both (B+C) markedly reduces viral entry. In Panels A and C, Calu-3 cells were transduced with nanoluciferase-expressing HIV particles pseudotyped with SARS-CoV-2 S WT, µS1/S2 or μS2’ for 72h and analyzed for nano-luciferase expression. Viral entry was expressed as fold increase over that given by bald particles (pseudovirions made in the absence of S). In panels **A** and **C** each dot represents a different experiment with median luciferase activity calculated from three biological replicates. Two to four experiments were performed for each cell type. Error bars indicate standard deviation (SD) from the mean.

Having observed the negative effect of BOS-318 on S processing by particle-producing cells, we asked whether pre-treating target cells with BOS-318 would also affect entry of SARS-CoV-2 pseudoparticles. In Calu-3 cells, where viral entry occurs primarily through fusion at the plasma membrane (46), we observed reduced viral entry by ∼3.8-fold for WT-S and ∼14-fold for μS2’-S (Fig. 6C). This emphasizes a significant contribution of Furin to the processing and priming of S at the plasma membrane of Calu-3 cells after ACE2 recognition. The fact that BOS-318 had a more pronounced effect on entry by μS2’-S-containing viral particles was not surprising given the very efficient S2’ processing of this mutant by Furin (Fig. 2D). Of note, viral entry by WT-S pseudoparticles was more affected by 100 μM Camostat compared to that by μS2’-S viral particles (Fig. 6C). Here, entry was reduced by ∼8.6-fold for WT-S and ∼3.2-fold for μS2’-S, suggesting that Furin plays a more prominent role in the entry of μS2’-S *versus* WT-S viral particles. Lastly, the combined pre-treatment of both BOS-318 and Camostat led to a complete block of viral entry, highlighting the importance of both Furin and TMPRSS2 in mediating viral fusion at the plasma membrane.

### Furin-like inhibitors decrease cell-to-cell fusion and syncytia formation

To assess whether BOS-inhibitors also affect cell-to-cell fusion, we developed a co-culture assay in which donor HeLa cells express HIV Tat and the fusogenic S-protein, while acceptor HeLa TZM-bl cells express ACE2 and Tat-driven luciferase (49) (Figs. 7A). As a proof-of-principle, we showed that when donor HeLa cells expressing HIV gp160 and Tat fused with acceptor cells expressing CD4 (SI-Fig. 6A, panel b), luciferase activity was increased compared to that observed in TZM-bl control cells co-cultured with donor Hela cells expressing only Tat (SI-Fig. 6B). The expression of S-protein alone in donor HeLa cells did not induce fusion with acceptor TZM-bl control cells (SI-Fig. 6B). However, ACE2 expression in TZM-bl allowed fusion with HeLa-expressing S-protein (SI-Fig. 6A, panel c; 6B) in a dose-dependent manner (SI-Fig. 6C), but no fusion was observed with μS1/S2-S (SI-Fig. 6A, panel d). Indeed, the linearity of our assay (correlation coefficient of 0.87) validated the use of luminescence as an indicator of cell-to-cell fusion (SI-Fig. 6C). Using this assay, we found that while donor cells expressing WT-S led to syncytia formation (Fig. 6B) and a >10-fold increased cell-to-cell fusion compared to control (empty vector V, no S) cells (Fig. 6C), donor cells expressing µS1/S2-S did not promote any cell fusion even in the presence of ACE2 (Figs. 6C, D). Thus, the µS1/S2-S phenocopies the effect of BOS-inhibitors on cell-to-cell-fusion (Fig. 6C), whereby absence of Furin-activity would not allow fusion and demonstrates a key role of S1/S2 cleavage in S-mediated cell-to-cell fusion. Consistent with this finding, we observed an almost complete loss of cell fusion when donor cells were treated with BOS-inhibitors or the PC-inhibitor decanoyl-RVKR-cmk (RVKR; Fig. 6C) (43), emphasizing the critical role of Furin-cleavage in promoting ACE2-dependent cell-to-cell fusion in the context of acceptor cells that do not endogenously express TMPRSS2, such as HeLa cells. This fusion assay also enabled the assessment of the effects of some worldwide-spreading S-protein variants of SARS-CoV-2, which seem to affect viral traits such as transmissibility, pathogenicity, host range, and antigenicity of the virus (50, 51). Among these, we selected mutants that modify the Pro at the P5 position of the S1/S2 site (Fig. 1A), i.e., the P681H and P681R associated with the α- and δ-variants, respectively (9). Our data showed that while the μS2’ mutant did not affect cell-to-cell fusion, the P681H and P681R mutants significantly enhanced it by ∼2-fold (Fig. 7D), in line with the higher transmissibility of the associated α- and δ-SARS-CoV-2 variants (50, 51) and increased cell-to-cell fusion (52).

**Figure 7:**
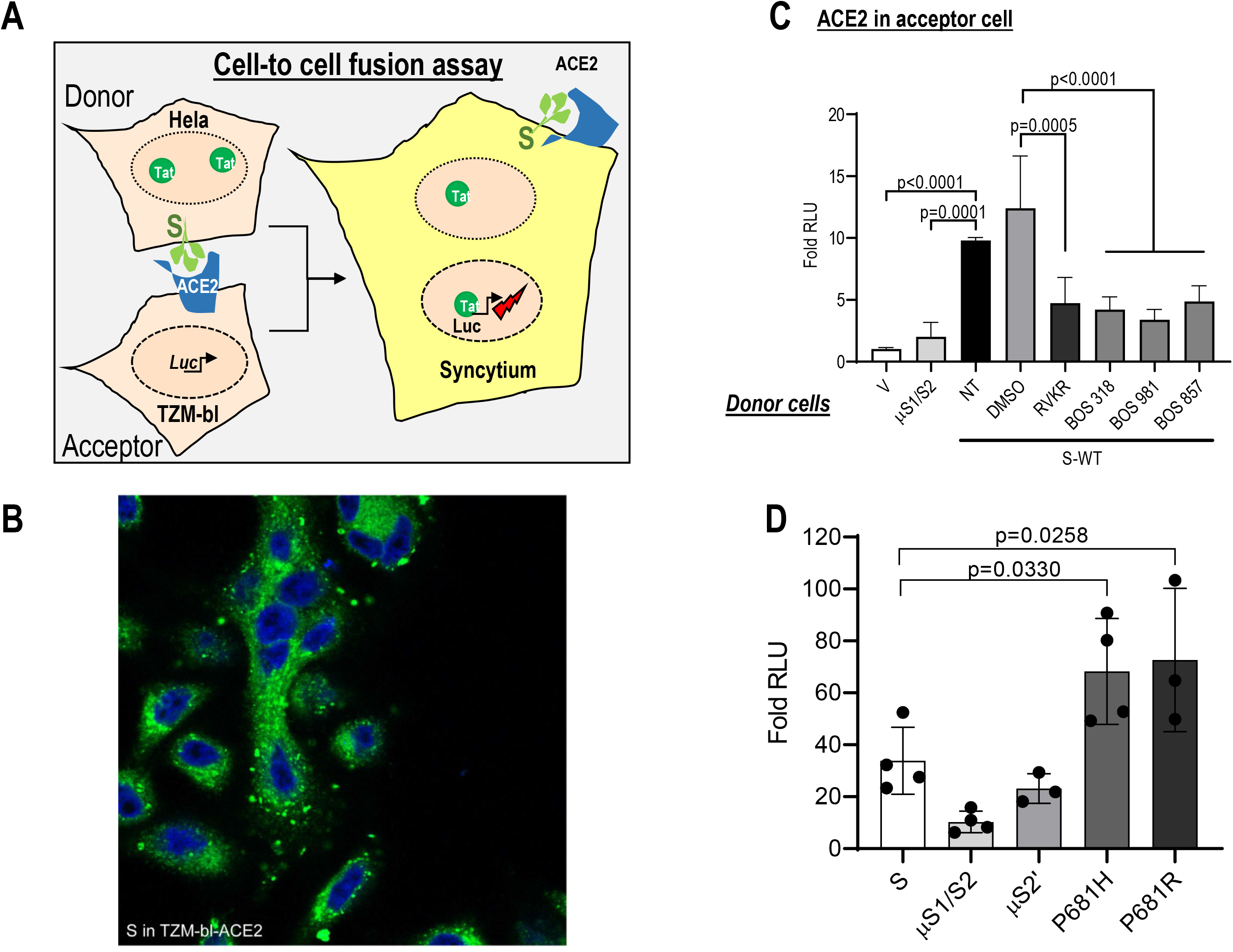
Spike-induced cell-to-cell fusion relies on Furin cleavage at S1/S2. (**A**) Cell-to-cell fusion between donor cells (HeLa) expressing the fusogenic SARS-CoV-2 spike protein along with the HIV trans-activator Tat, and acceptor cells (TZM-bl) that express ACE2. Upon fusion, Tat is transferred from donor to acceptor cells, thereby inducing luciferase expression. (**B**) Cell-to-cell fusion was evaluated using confocal microscopy. A representative immunocytochemistry of Hela cells transfected with a vector expressing SARS-CoV-2 spike co-cultured with TZM-bl cells for 18h. The number of syncytia (multiple nuclei) was examined using CellMask™ to probe for the plasma membrane and Dapi to stain the nuclei. (**C**) Donor cells were transfected with vectors expressing either no protein (empty vector, V), μS1/S2, or WT-spike (S) in the absence (NT) or presence of vehicle (DMSO) or with the Furin-inhibitors BOS-318, BOS-981, BOS-857 (300 nM) or RVKR (10 μM). Acceptor cells were transfected with a vector expressing ACE2. After 48h, donor and acceptor cells were co-cultured for 18h. Relative luminescence units (RLU) were normalized to the V value arbitrarily set to 1. Data are presented as mean values ± SD (n=3), One-Way ANOVA, Dunn-Sidàk multiple comparison test. (**D**) Donor HeLa cells expressing WT-S or its indicated mutants and variants were co-cultured with acceptor TZM-bl cells expressing ACE2. The extent of fusion is represented as a ratio between the RLU measured for each condition and that of donor cells expressing empty vector. The bar graph represents the average of 3 experiments performed in triplicates. Data are presented as mean values ± SEM (n=3), One-way Anova Bonferroni multiple comparison test. Two-Way ANOVA, Dunn-Sidàk multiple comparison test.

### TMPRSS2 promotes cell-to-cell fusion in the absence of Furin-mediated cleavage at S1/S2

Having shown the critical role of Furin-cleavage in promoting ACE2-dependent cell-to-cell fusion in HeLa cells that do not express TMPRSS2, we next examined the importance of TMPRSS2 in cell-to-cell fusion given its significant role in viral entry and replication (Figs. 5D and 6C). Thus, we analyzed fusion between donor cells expressing WT-S or μS1/S2-S (double tagged S-protein: HA at the N-terminus and V5 at the C-terminus) and acceptor cells expressing ACE2 in the absence or the presence of TMPRSS2 (Fig. 8). As shown, in the presence of ACE2, TMPRSS2 reduced fusion by ∼2.1-fold in the case of WT-S but increased it by ∼2.4-fold in the case of μS1/S2-S (Fig. 8A). Under the same conditions, WB analyses showed that for WT-S, the TMPRSS2 expression decreased by ∼4-fold (3.8 *versus* 0.9) the relative levels of S2 and modestly increased by ∼1.5-fold (0.6 *versus* 0.9) the relative levels of S2’. In contrast, for μS1/S2-S, expression of TMPRSS2 increased S2’ levels by ∼7-fold (0.1 *versus* 0.7) without any detectable change in S2 (0.6 *versus* 0.5) (Fig. 8B). Therefore, in the presence of ACE2, the levels of S2 correlate with cell-to-cell fusion in the case of WT-S that is well cleaved at S1/S2 by Furin. In contrast, fusion correlates best with S2’ levels, when cleavage at S1/S2 is limited (μS1/S2-S). Interestingly, secretion of the HA-tagged, N-terminal S1 subunit (aa 14-685; Fig. 1A) was more pronounced for WT-S compared to μS1/S2-S (Fig. 8B). Yet, in both cases, the extent of S1 release was not modulated by the presence of TMPRSS2, suggesting that TMPRSS2-mediated enhanced fusion of µS1/S2 was not a consequence of altered S1/S2 cleavage (Fig. 8B).

**Figure 8:**
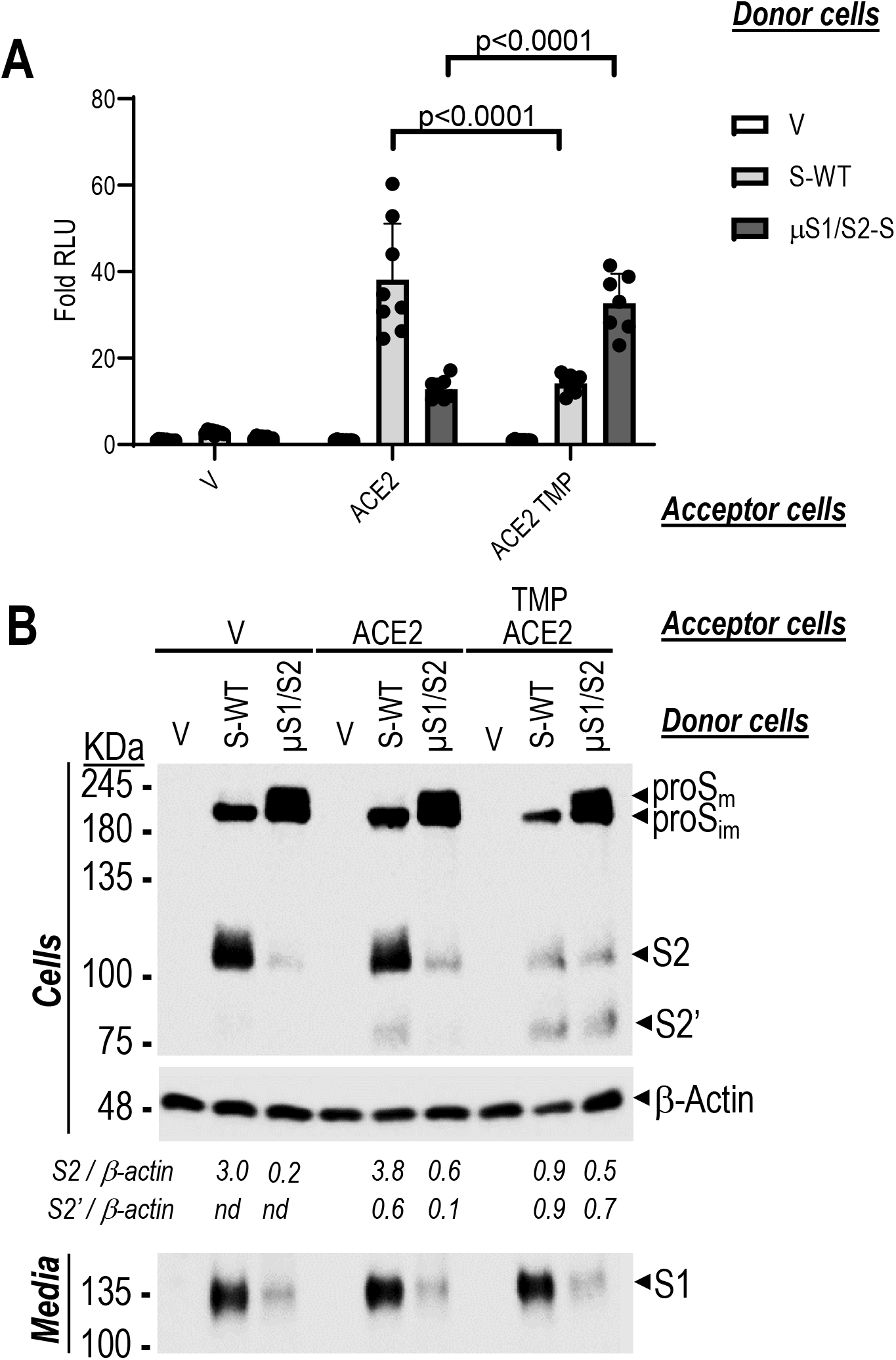
Spike-glycoprotein processing by Furin and TMPRSS2 in a co-culture system and their role in cell-to-cell fusion. Donor HeLa cells expressing empty vector (V), WT-S-HA or μS1/S2-HA were co-cultured with acceptor TZM-bl cells expressing V, ACE2, or ACE2 + TMPRSS2 (TMP). From the same experiment, cell-to-cell fusion (**A**) was assessed in parallel by Western blotting of spike-glycoproteins in cells and media using an anti-V5 mAb (**B**). Secreted forms of spike protein (S1) in the media were detected with anti HA-HRP upon immunoprecipitation with anti-HA agarose. The bar graph shows one representative fusion assay done un duplicate. The corresponding Western-blot is representative of three independent experiments. Values of S2 and S2’ relative to β-actin are shown (nd = too low or not detected).

### Mechanism by which TMPRSS2 promotes cell-to-cell fusion

To assess whether TMPRSS2 promoted cell-to-cell fusion by enhancing S2’ cleavage in the complete absence of S1/S2 priming, we generated a new S-derivative lacking all Arg at S1/S2 (Fig. 1), namely μAS1/S2 (AAAA_685_)-S, and hence different from μS1/S2 (Fig. 1C; ARAA mutant), it would not be cleaved by TMPRSS2 or Furin. Cell-to cell fusion was assessed following incubation of cells expressing WT-S or μAS1/S2-S with acceptor cells expressing ACE2 and/or TMPRSS2. In this context, TMPRSS2 reduced fusion of WT-S by ∼2.6-fold and enhanced that of µAS1/S2-S by ∼3.1-fold (Fig. 9A), as previously observed with the μS1/S2 mutant (Fig. 8A). Camostat completely restored TMPRSS2-reduced fusion with WT-S and largely attenuated the TMPRSS2-enhanced fusion with µAS1/S2 (Fig. 9A). This revealed that in the presence of ACE2, TMPRSS2 could enhance fusion in absence of S1/S2 priming. We next assessed whether this effect could be related to the differential processing of the S protein and/or to ACE2 receptor shedding by TMPRSS2. The WB of cell lysates shows that treatment of acceptor cells with Camostat eliminated the TMPRSS2-induced reduction in S2 observed with WT-S (Fig. 9B), yet it did not significantly alter S2’ levels either with WT-S or µAS1/S2 (Figs. 9B, C). In contrast, the WB of ACE2 reveal that TMPRSS2 strongly reduced the levels of mature membrane-bound ACE2 migrating at ∼120 KDa, without affecting the lower molecular weight protein which correspond to an immature ER-retained form of ACE2 (ACE2_im_, Figs. 9B, C) that is sensitive to endoH digestion (*not shown*). These results show that ACE2 is shed by TMPRSS2, as proposed in earlier studies (33). As expected, we observed that ACE2 shedding by TMPRSS2 is almost completely blocked by Camostat (compare ACE2+TMP lanes DMSO to CAM) (Figs. 9B and C).

**Figure 9:**
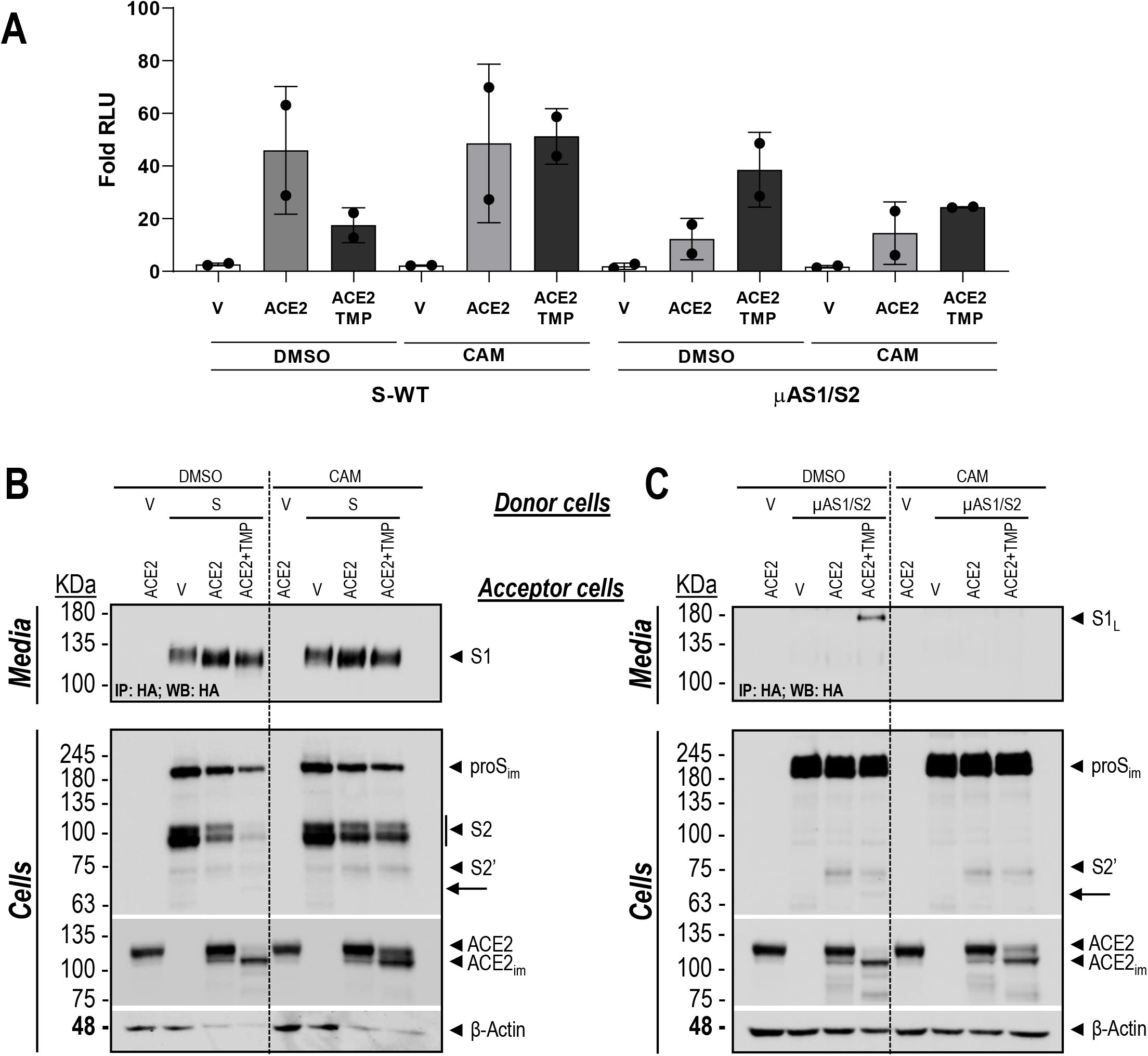
Exogenous TMPRSS2-generated shedding of ACE2 differentially regulates S-induced fusion at the plasma membrane of WT-S *versus* µAS1/S2. Donor HeLa cells expressing double tagged (N-terminal HA-tag; C-terminal V5-tag) spike-glycoprotein WT (S) or its S1/S2 mutant (µAS1/S2) were co-cultured with acceptor TZM-bl cells expressing (empty vector, V), ACE2, or ACE2 + TMPRSS2 (TMP) and treated with DMSO (vehicle control) or Camostat (120 μM). Within the same experiment, cell-to-cell fusion (**A**) was assessed in parallel with spike processing in cells and media by Western-blot (**B, C**). (**A**) The extent of fusion is represented as a ratio between the RLU measured for each condition and that of donor cells expressing V. The bar graph represents the average of 2 experiments performed in triplicates. (**B, C**). Western blot analyses of media and cell extracts of the co-cultured cells with donor cells overexpressing double tagged (N-terminal HA-tag; C-terminal V5-tag) spike-glycoprotein, WT (S) (**B**) or µAS1/S2 (**C**). The arrow points to a putative degradation product of S2’, that is absent in presence of Camostat. Media were subjected to immunoprecipitation with anti-HA agarose for the secreted forms of spike protein (S1, S1_L_) followed by Western blotting with anti HA-HRP. In the cell extracts Spike-glycoproteins and ACE2 were immunoblotted with anti-V5 mAb and a polyclonal ACE2 antibody, respectively. The data are representative of three independent experiments.

In contrast, the WB anti ACE2 reveal that TMPRSS2 strongly reduced the levels of mature membrane-bound ACE2 migrating at ∼120 KDa, without affecting the lower molecular weight protein which correspond to an immature ER-retained form of ACE2 (ACE2_im_, Figs. 9B, C) that is sensitive to endoH digestion (*not shown*). These results suggest that ACE2 is shed due to TMPRSS2 cleavage as proposed in earlier studies (33). As expected, we observed that this effect of TMPRSS2 on ACE2 processing is almost completely blocked by Camostat (compare ACE2+TMP lanes DMSO to CAM) (Figs. 9B and C). Finally, by analysing the release of S1 in the cell culture media we observed that TMPRSS2-induced release into the media of a longer ∼175 kDa fragment from µAS1/S2, referred hereafter as S1_L_ (Fig. 9C), which is not detected in presence of Camostat. The molecular weight of the S1_L_ indicates that it corresponds to a TMPRSS2 cleavage of µAS1/S2-S at S2’ (Fig. 9B).

These results suggest that TMPRSS2 modulation of fusion is complex, as TMPRSS2 alters both the integrity of ACE2 receptor, but also participates in S-processing as attested by the release of S1_L_ which corresponds to the N-terminal cleavage product of µAS1/S2-S at S2’. Interestingly, we did not detect an associated increase in S2’ levels, suggesting that after cleavage the fusion process occurs and the S2 protein is subsequently degraded, as suggested by a small degradation product seen on WB (arrow in Figs. 9B, C). We surmise that in the presence of ACE2, TMPRSS2 facilitates the secretion of the N-terminal fragment S1_L_ generated by cleavage at S2’.

Our next goal was to further decipher the contribution of ACE2 cleavage by TMPRSS2 and the seemingly opposite effects of TMPRSS2 on fusion activity with WT-S and µAS1/S2-S (and µS1/S2-S). Hence, since TMPRSS2 cleaves at single Arg and Lys residues (53), we tested the effect of multiple Arg/Lys to Ala mutants in segments close to the transmembrane domain of ACE2 (C0 + C4 in Fig. 10A), which were previously proposed to limit the TMPRSS2-induced shedding (54). Accordingly, WB analyses (Fig. 10B) revealed that TMPRSS2, but not its Ser* active site S441A mutant (μTMPR, Fig. 10B; SI-Fig. 7) sheds the membrane-bound ∼120 kDa WT ACE2, releasing a major ∼95 kDa and a minor ∼80 kDa form into the media. In contrast, TMPRSS2 primarily generated the ∼80 kDa fragment from the ACE2 (C0+C4) mutant. It should be mentioned that in the absence of TMPRSS2 the extent of fusion for both WT-S and µAS1/S2-S was comparable between ACE2 (WT) and ACE2 (C0+C4). Interestingly, in the context of the ACE2 (C0+C4) receptor mutant, we found that TMPRSS2 no longer modulates the fusion of both WT-S and µAS1/S2-S (Fig. 10C). Importantly, and different from WT ACE2 (Figs. 8, 9), TMPRSS2 did not reduce the levels of S2 when ACE2 (C0+C4) was used (Fig. 10D), nor did it reduce fusion (Fig. 10C). Additionally, the TMPRSS2-enhanced secretion of the ∼ 175 kDa S1_L_ fragment in the absence of S1/S2 cleavage (Fig. 9C; SI-Fig. 7), was no longer detected in the media when ACE2 (C0+C4) was expressed on acceptor cells (Fig. 10D), correlating with the lack of fusion of µAS1/S2-S (Fig. 10C). Altogether, these results suggest for the first time that in the absence of Furin-mediated priming of S, TMPRSS2 promotes cell-to-cell fusion by generating a ∼95 kDa sACE2 that likely binds the S1_L_ cap (55) and favors its release.

**Figure 10:**
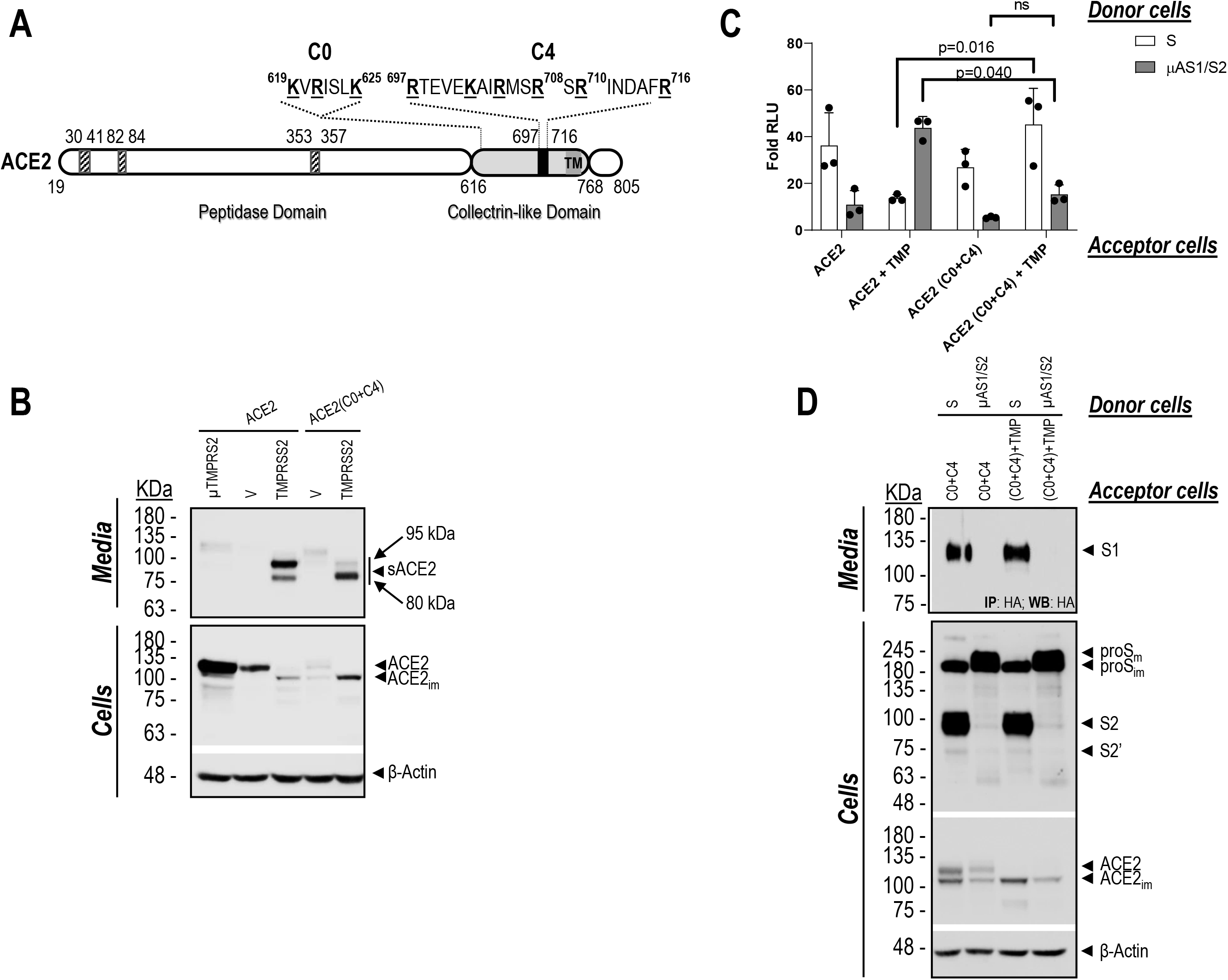
The C-terminal collectrin-like domain of ACE2 may be critical for the regulation of cell-to-cell fusion of spike-glycoprotein when exogenous TMPRSS2 is present. **(A)** Schematic representation of the primary structure of human ACE2 with emphasis on the C-terminal collectrin-like domain (aa 616-768, light gray), TMPRSS2 cleavage region (aa 697-716, black) and the polybasic amino-acid segments in which K/R were mutated to A (C0 and C4) (amino acids underlined and in bold). Also shown are the peptidase domain (aa 19-615, white) containing the regions involved in the interaction with the spike SARS-CoV protein (hatched) and transmembrane domain (TM). (**B**) HeLa cells were co-transfected with ACE2, WT (ACE2) or its mutant ACE2 (C0+C4), and TMPRSS2, WT (TMPRSS2) or its S441A active-site mutant (µTMPRSS2), or empty vector (V). Media and cell extracts were analyzed by western blotting for shed ACE2 (sACE2) and ACE2, respectively. The migration positions of the ∼95 kDa and ∼80 kDa sACE2 are emphasized. (**C**) Donor HeLa cells expressing WT-S-HA or μAS1/S2-HA were co-cultured with acceptor TZM-bl cells expressing ACE2, WT (ACE2) or its mutant ACE2 (C0+C4) in presence or absence of TMPRSS2. From the same experiment, cell-to-cell fusion was assessed (**C**), in parallel with WB analyses of cells and media (**D**). The extent of fusion is represented as a ratio between the RLU measured for each condition and that of donor cells expressing an empty vector. The bar graph represents the average of 3 experiments performed in triplicates. Data are presented as mean values ± SD (n=3), One-way Anova Turkey’s multiple comparison test. (**D**) Co-culture media were subjected to immunoprecipitation with anti-HA agarose for the secreted forms of spike protein (S1) followed by western blot with anti HA-HRP. Spike-glycoproteins in the cell extracts were immunoblotted with anti-V5 mAb. The Western blot data are representative of three independent experiments.

## DISCUSSION

Furin and TMPRSS2 have both been proposed to process the SARS-CoV-2 spike protein and promote viral entry and infection. In this study, we functionally characterize the role of Furin and TMPRSS2 and show their complementary role in SARS-CoV-2 mechanism of entry and infection.

Consistent with previous data (18, 33, 56, 57), processing at S1/S2 is required for optimal viral entry in Calu-3 lung epithelial cells. Indeed, mutation of the arginines of the FCS or blocking S1/S2 cleavage by a series of novel Furin-inhibitors effectively reduced SARS-CoV-2 entry in Calu-3 cells. In the context of cell-to-cell fusion, mutations of the FCS impaired syncytia formation in HeLa cells in the absence of TMPRSS2 in acceptor cells, and Furin-inhibitors phenocopied the effect of these S1/S2 mutations. These results highlight the importance of Furin in promoting S-mediated cell-to-cell fusion. In contrast, in HEK293T cells, where SARS-CoV-2 likely enters *via* endocytosis in acidic endosomes, mutations of the FCS enhanced entry (Fig. 6), suggesting that cleavage at S1/S2 by Furin was not required for efficient S-processing by endosomal proteases (e.g., cathepsins) (58, 59). The pH-dependent entry pathway is sensitive to drugs increasing the pH of endosomal compartments, e.g., hydroxy-chloroquine (47). Nevertheless, this drug failed to show significant improvement in COVID-19 patients (60, 61) or infected animals (62), supporting the notion that the virus can effectively utilize alternative entry pathways that are critical for virus entry, transmission and pathogenicity. The fact that Furin-cleavage at S1/S2 is conserved in SARS-CoV-2 isolated from COVID-19 patients and that ΔPRRA viruses are poorly infectious in hamsters (60), suggest that blocking viral entry through the pH-independent pathway is a viable approach towards thwarting SARS-CoV-2 dissemination. In this context, the non-toxic, small molecules BOS-inhibitors that were analyzed in this study, which can be delivered orally or by inhalation, deserve consideration as potential antivirals against acute SARS-CoV-2 infection. As observed in adult animal models, short-term inhibition of Furin would not cause severe side effects, despite the many physiological functions of this enzyme (20).

Our results also demonstrate that Furin can also process S at S2’, a site that we map by proteomics to be at **<u>KR</u>**_815_↓SF. The latter S2’ cleavage site was confirmed by introducing an optimized S2’ site (μS2’, KRRKR_815_↓SF) that was efficiently cleaved by Furin, yielding a protein fully competent for fusion in pseudotyped experiments. Importantly, we showed that the S2’ processing is strongly increased when ACE2 receptor is expressed. Our results suggest that the recognition of the ACE2 receptor by the spîke protein induces a conformational change of the S2 domain and enhances cleavage of S2’ site by cellular proteases such as Furin.

In contrast TMPRSS2 appears to cleave synthetic peptides encompassing the S1/S2 and S2’ cleavage sites and proS protein in Hela cells less efficiently compared to Furin. TMPRSS2 has been reported to enhance the infectivity of SARS-CoV-1 and MERS-CoV *via* cleavages of the S-protein (53, 63), as reviewed in (4, 15, 23). The S1/S2 site VSLL**<u>R</u>**_667_↓ST of SARS-CoV-1 contains an Arg↓Ser cleavage motif that is propitious to cleavage by TMPRSS2 (53) but not by Furin. However, S1 production was minimal and fusion was observed only when donor cells expressing S-protein were co-cultured with acceptor cells expressing ACE2 and TMPRSS2 (53). In SARS-CoV-2 the S1 released, presumably following cleavage of S at S1/S2 and ACE2 binding, is much reduced with µS1/S2-S compared to WT-S, indicating that Furin-mediated cleavage at S1/S2 promotes a more efficient release of S1 (Fig. 8B). Additionally, in a co-culture system the relative levels of soluble S1 fragment were not affected by the presence or absence of TMPRSS2 for WT and µS1/S2 (Fig. 8B). This suggests that the differential effect of TMPRSS2 on cell-to-cell fusion could not be attributed to its direct cleavage activity on S. Rather, we assert that TMPRSS2 primarily modulates cell-to-cell fusion in part *via* shedding of ACE2. In the presence of TMPRSS2 the extent of fusion would depend on a combination of reduced levels of full-length cell surface ACE2 and increased levels of sACE2 that could act as a decoy to inhibit fusion (64). In that context, a human recombinant sACE2 that includes the collectrin domain but lacks the TM domain (hrsACE2, aa 1-740) (see Fig. 10A) effectively blocked SARS-CoV-2 infection of Vero E6 cells (55). Indeed, our cell-to-cell fusion assay tested in co-culture revealed that the ∼95 kDa sACE2 primarily produced by TMPRSS2 cleavage of ACE2 resulted in lower levels of cellular ACE2 (Figs. 9B, 10B) and in impaired WT-S induced fusion (Figs. 9A, 10C). In contrast, the mutant ACE2 (C0+ C4) was no longer sensitive to TMPRSS2 impairment of WT-S-induced fusion, suggesting that the ∼80 kDa form of sACE2 primarily produced by TMPRSS2 cleavage of ACE2 (C0+C4), likely upstream of Lys_619_, may no longer be able to inhibit WT-S-fusion (Fig. 10C), but may still bind S1 as shown by cryoEM studies that used a short ACE2 ectodomain (aa 19-615) (40). Altogether, these results suggest that the C-terminal collectrin-like domain (65, 66) of ACE2 (aa 616-768) (Fig. 10A), which is lost in the ∼80 kDa form, may be critical for the ability sACE2 to inhibit cell-to-cell fusion of WT-S. Since Arg_710_ and Arg_716_ within the collectrin-like domain (Fig. 10A) have been reported to be implicated in ACE2 dimerization (66), their Ala mutation in the ACE2 (C0+C4) mutant should significantly impair ACE2 dimerization. Whether dimerization of ACE2 and the ∼95 kDa sACE2 (Fig. 10B) are needed for efficient inhibition of WT-S-induced fusion (64) is yet to be confirmed.

In the context of endogenous expression of ACE2 and TMPRSS2, e.g., in Calu-3 cells (67, 68), our data show that Camostat significantly reduces SARS-CoV-2 infectivity (Fig. 6C), revealing that a TMPRSS2-like activity favors viral entry. Similarly, in the presence of BOS-inhibitors or absence of Furin cleavage at S1/S2, TMPRSS2 enhances cell-to-cell-fusion (Figs. 8A, 9A). HeLa cells expressing WT-S in donor cells and both ACE2 and TMPRSS2 in acceptor cells, exhibited reduced cell-to-cell fusion, a process inhibited by Camostat. This suggests that the relative contribution of TMPRSS2 to the shedding of ACE2 into sACE2 and cleavage of S-protein at S2’ is cell-type dependent. Future experiments may unravel the underlying mechanism that would explain the difference between the HeLa and Calu-3 cells. Our study highlights a complex dynamic between spike, ACE2, Furin and TMPRSS2 (Fig. 11) and points to a potential role of TMPRSS2, which in the absence of S1/S2 processing (e.g., μAS1/S2 or in presence of BOS-inhibitors) can by shedding ACE2 facilitate S2’ cleavage and cell-to-cell fusion.

**Figure 11:**
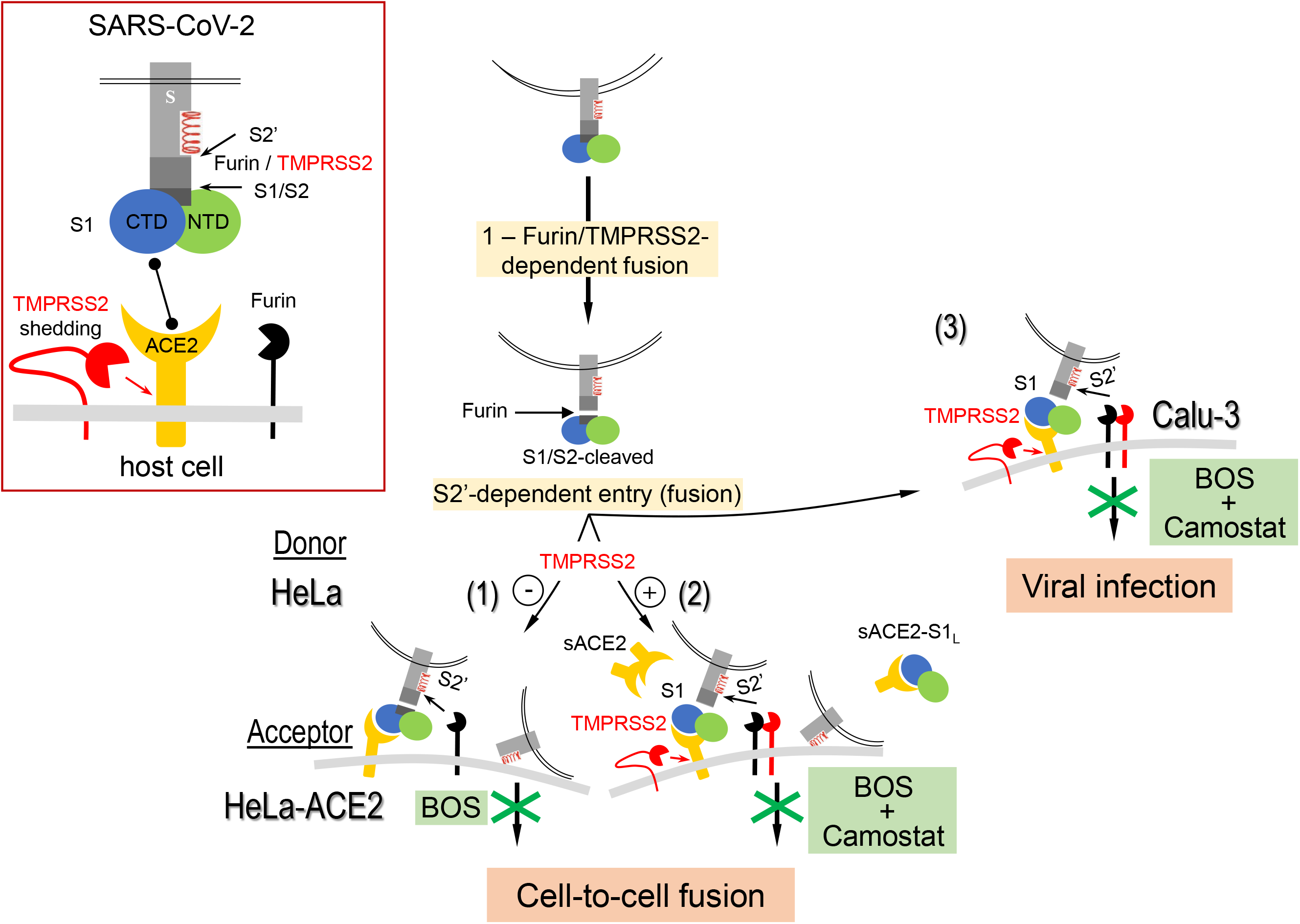
Proposed model for the processing of S-protein and its blockade by Furin and TMPRSS2 inhibitors. <u>Boxed left panel</u>: schematic representation of the S-glycoprotein domains of SARS-CoV-2, including the N-terminal (NTD) and C-terminal (CTD) domains of S1, the Furin-S1/S2 and the Furin/TMPRSS2-S2’ processing sites as well as the fusogenic α-helix that follows S2’. Binding of the RBD domain of S1 to the membrane associated ACE2 in target cells, and the cell surface expression of TMPRSS2 and Furin are also schematized. <u>Right panels</u>: (**1**) BOS-inhibitors (or μS1/S2 mutant) completely prevent cell-to-cell fusion of donor Hela cells expressing S-glycoprotein with acceptor HeLa-ACE2 cells, which lack endogenous TMPRSS2. In this context, this reveals that Furin is a major processing enzyme cleaving at S1/S2 and generating S2’. (**2**) In acceptor HeLa cells expressing TMPRSS2 (+), maximal prevention of cell-to-cell fusion can be achieved by a combination of Furin (BOS, phenocopying the μS1/S2 or μAS1/S2 mutants) and TMPRSS2 (Camostat) inhibitors blocks S2’ production, ACE2-shedding (sACE2) and the separation of sACE2-S1_L_ complex from S2. (**3**) Optimal blockade of SARS-CoV-2 infection of Calu-3 cells, which express endogenously both Furin and TMPRSS2, is also achieved by a combination of Furin (BOS) and TMPRSS2 (Camostat) inhibitors.

The human airway epithelium is an important site of early SARS-CoV-2 infection (18, 24, 69). The virus can then disseminate to other tissues/cells such as gut, liver, endothelial cells and macrophages where ACE2, Furin and TMPRSS2 are co-expressed (70). While Furin is mostly found in the TGN, it is also present in endosomes and on apical/basolateral plasma membranes in polarized cells such as those of the lung, small intestine and kidney (71). In contrast, TMPRSS2 is mostly found at the apical membrane of secretory epithelia (72), suggesting that both enzymes would be poised to process the S-protein on the apical side. Their relative abundance may be an important factor governing which of the two might cleave S at the S2’ cleavage site. The complementarity and interchangeability of these different proteases, together with those in endosomes, likely allows SARS-CoV-2 to exhibit a wider tropism compared to SARS-CoV-1 (73). In this context, we note that randomized placebo controlled clinical trials using an orally administered TMPRSS2 inhibitor Camostat mesilate three times a day for treatment of COVID-19 patients did not show improvement in outcomes (74). We believe that the maximal benefit of TMPRSS2 inhibition could be achieved when Furin activity is also inhibited, since our results showed that TMPRSS2 activity is preponderant in the context of absence of cleavage at S1/S2. Furthermore, the intranasal delivery of both agents would be expected to have less side effects and be more effective, as it would directly target the airway epithelia of the nose and the lungs, the major sites of SARS-CoV-2 entry.

Altogether, our results strongly support the notion that a combination of BOS-and selective TMPRSS2-inhibitors would provide a more effective blockade against SARS-CoV-2 infection (Figs. 5D, 11). It would now be interesting to validate *in vivo* whether a combination of a BOS-inhibitor and a more potent/selective TMPRSS2-inhibitor (75) together with hydroxy-chloroquine (56, 76) would synergize the antiviral effect of these entry inhibitors.

The availability for worldwide distribution of various SARS-CoV-2 vaccines that inhibit the accessibility of the RBD of S-protein to ACE2 (https://www.raps.org/news-and-articles/news-articles/2020/3/covid-19-vaccine-tracker) represents a major advance to limit the spread of SARS-CoV-2 infections. However, it is still not known with certainty whether they will be effective in patients with impaired immune systems, and whether they will confer a persistent protection against new variants of SARS-CoV-2. If the protective effect of the vaccination remains incomplete, effective antiviral drugs are still needed and could help with early diagnosis of the disease. Ultimately, in case of new emerging coronavirus pandemics (77), the availability of such treatments would constitute a powerful anti-viral arsenal to be used in pandemic preparedness.

## MATERIALS AND METHODS

### Enzymatic PC-inhibition by BOS-inhibitors

#### Biochemical assay

The proprotein convertases Furin (108-574-Tev-Flag-6His), PC5A (PCSK5; 115-63-Tev-Flag-6His), PACE4 (PCSK6; 150-693-Tev-Flag-6His), and PC7 (PCSK7; 142-634-Tev-Flag-6His) enzymes were purified from BacMam transduced CHO cells. Reactions were performed in black 384-well polystyrene low volume plates (Greiner) at a final volume of 10 µL. BOS-inhibitors (BOS-318, BOS-857 and BOS-981) were dissolved in DMSO (1 mM) and serially diluted 1 to 3 with DMSO through eleven dilutions to provide a final compound concentration range from 0.00017 to 10 μM. 0.05 μl of each concentration was transferred to the corresponding well of an assay plate, and then 5 μl of enzyme (Furin, PCSK5, PCSK6, and PCSK7) in assay buffer (100 mM HEPES pH7.5, 1 mM CaCl_2_ and 0.005% Triton X-100) was added using a Multidrop Combi (Thermo) to the compound plates to give a final protein concentration of 0.02, 0.5, 2.5, and 1.0 nM respectively. The plates were mixed by inversion and following a 30 min preincubation of enzyme with compound at room temperature (∼22°C), the substrate FAM-QRVRRAVGIDK-TAMRA (AnaSpec # 808143, 5 μl of a 1, 0.25, 0.20, and 0.5 μM solution in assay buffer for Furin, PCSK5, PCSK6, and PCSK7 respectively) was added using a Multidrop Combi. The plates were centrifuged at 500 x *g* for 1 minute and incubated at room temperature for two hours. Enzyme inhibition was then quantified using an Envision instrument (PerkinElmer). Data were normalized to maximal inhibition determined by 1 μM decanoyl-Arg-Val-Lys-Arg-chloromethylketone (Calbiochem #344930).

#### Golgi imaging assay

This assay uses an image-based platform to evaluate the intracellular activity of Furin inhibitors. Reactions were performed in black 384-well, tissue culture-treated, clear bottom plates (Greiner). Compounds dissolved in DMSO (1.0 mM) were serially three-fold diluted to give a final compound concentration range from 0.00017 to 10 μM. Analyses were initiated by the addition of U2OS cells simultaneously transduced with a BacMam-delivered construct containing a Golgi-targeting sequence followed by a 12-amino acid Furin/PCSK cleavage site from Bone Morphogenic Protein 10 (BMP10) and then GFP at the C terminus. The dibasic Furin cleavage site sequence was flanked by glycine rich linkers (GalNAc-T2-GGGGS-DSTARIRRNAKG-GGGGS-GFP). Briefly, frozen cells are thawed in assay media (Dulbecco’s Modified Eagles Medium Nutritional Mixture F-12 (Ham) without phenol red containing 5% FBS) and diluted to deliver 6000 cells/well (50 μl) to the plate using a Multidrop Combi (Thermo). After a 24-hour incubation period at 37°C, the cells are stained with Cell Mask Deep Red, fixed in paraformaldehyde and the nuclei stained using Ho33342. The Golgi-targeted GFP forms bright punctate clusters within the cell. In the absence of a Furin/PCSK inhibitor, the endogenous protease cleaves GFP from its N-acetylgalactosaminyltransferase-2 Golgi tether, releasing GFP into the Golgi lumen where fluorescence was diluted below the threshold of assay sensitivity. In the presence of a cell permeable Furin/PCSK inhibitor, GFP fluorescence increases as intra-Golgi protease activity was reduced. Cellular GFP intensity was determined by image-based acquisition (Incell 2200, Perkin Elmer) at 40x magnification with 4 fields measured per well. Multi-scale top hat segmentation was used to identify the GFP-tagged puncta and to quantitate the average fluorescence of all puncta on a per cell basis. Cellular toxicity was determined in parallel.

#### Furin and TMPRSS2 fluorogenic assays

Recombinant Furin was purchased from BioLegend (#719406), human recombinant TRMPSS2 from Cliniscience (ref LS-G57269-100), and the DABCYLGlu-EDANS labelled peptides encompassing the different cleavage sites (SI Table 1) were purchased from Genscript. Reactions were performed at room temperature in black 384-well polystyrene low volume plates (CELLSTAR-Greiner Bio-One # 784476) at a final volume of 15 µL. The fluorescent peptides were used at 5 µM and the reactions were performed in 50 mM Tris buffer (pH 6.5 or 7.5), 0.2% Triton X-100, 1mM CaCl_2_ and Furin was added at a final concentration of 0.2 to 100 nM. BOS-inhibitors (BOS-318, BOS-857 and BOS-981) were dissolved in DMSO (1 mM) and serially diluted 1 to 2 with DMSO to provide a final compound concentration range from 50 μM to 0.01 nM with 5% DMSO in the enzymatic assay. For TMPRSS2, the fluorescent peptides were used at 5 µM and the reactions were performed in 50 mM Tris buffer (pH 8), 150 mM NaCl and TMPRSS2 was added at final concentrations of 50 nM. Cleavage of the synthetic peptides was quantitated by determining the increase of EDANS (493 nM) fluorescence following release of the DABCYL quencher, which was excited at 335 nM using a Safire 2 Tecan fluorimeter. The fluorescence was followed during 90 min, and the enzymatic activity was deduced by measurement of the increase of fluorescence during the linear phase of the reaction. Each reaction was performed in triplicate and the standard deviation was calculated using Excel-ecart type function (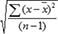).

### Plasmids

Single tagged (C-terminal V5 tag) or double tagged (N-terminal HA tag and C-terminal V5 tag) spike-glycoprotein of SARS-CoV-2 (optimized sequence) and its mutants were cloned into the pIRES2-EGFP vector. Site-directed mutagenesis was achieved using a Quick-Change kit (Stratagene, CA) according to the manufacturer’s instructions. The plasmids pCI-NEO-hACE2 received from DW Lambert (University of Leeds) and pIRES-NEO3-hTMPRSS2 from P Jolicoeur (IRCM). The ΔEnv Vpr Luciferase Reporter Vector (pNL4-3.Luc.R-E-) was obtained from Dr. Nathaniel Landau through the NIH AIDS Reagent Program whereas the pHIV-1NL4-3 ΔEnv-NanoLuc construct was a kind gift from Dr. P Bieniasz. Plasmids encoding VSV-G, as HIV-1 Env and tat were previously described (78, 79).

### Cell culture and transfection

Monolayers of HeLa, HEK293T, HEK293T17, Vero E6 and Calu-3 cells were cultured in 5% CO_2_ at 37°C in Dulbecco’s modified Eagle’s medium (DMEM; Wisent) supplemented with 10% (v/v) fetal bovine serum (FBS; Wisent). HEK293T-ACE2(80), a generous gift from Dr. Paul Bieniasz, were maintained in DMEM containing 10% FBS, 1% nonessential amino acids (NEAA) and 50 µg/ml blasticidin (Invivogen). The cells were transfected with JetPrime transfection reagent according to the manufacturer’s instructions (Polyplus transfection, New York, USA). At 24h post transfection, the culture media were changed to serum-free and cells incubated for an additional 24h. To establish the HeLa cells stably express human ACE2, transfected cells were selected using media containing 500 μg/ml of neomycin (G418, Wisent).

For knockdown of Furin in HeLa cells, an optimized set of 4 small interfering RNAs (siRNAs; SMARTPool) targeted against human Furin were purchased from Horizon Discoveries (Perkin Elmer, Lafayette, LA, USA) and transfections of HeLa cells were carried out using INTERFERin (PolyPlus) as recommended by the manufacturer.

To generate HIV particles pseudotyped with SARS-CoV-2 S, 293T17 cells (600,000 cells plated in a 6-well vessel) were transfected with 1 µg pNL4-3 Luc.R-E- (or pHIV-1NLΔEnv-NanoLuc) in the presence or absence of 0.3 µg pIR-2019-nCoV-S V5 plasmids using Lipofectamine-3000 (Life Technologies). In certain experiments, 293T17 cells were treated with BOS-inhibitors at 6 h post transfection. Pseudovirions expressing the nano- or firefly-luciferase were collected at 24 h or 48 h post transfection, respectively. Viral supernatants were clarified by centrifugation at 300 x *g*, passed through a 0.45-μm pore-size polyvinylidene fluoride (PVDF; Millipore) syringe filter (Millipore; SLGVR33RS), and aliquots frozen at −80°C. For WB analysis of purified pseudovirions, viral supernatants were concentrated by ultracentrifugation on a 20% sucrose cushion for 3h at 35,000 RPM; Beckman Coulter OPTIMA XE; Ti70.1 rotor). HIV particles lacking the SARS-CoV-2 S glycoprotein served as a negative control in all experiments.

### Cell viability assay using MTT

Cells, seeded in a 96-well plate, the day before, at 10,000 (HEK-293T and Vero E6) or 50,000 (Calu-3) cells, were treated with serial 10-fold dilutions of BOS-inhibitors for up to 48h. Cells treated with vehicle alone were used as negative control. MTT was subsequently added to the medium (final concentration: 2.5 mg/ml) and cells were further incubated for 4h at 37 ^0^C. After removal of the culture media, DMSO was added and absorbance read at 595 nm using a microplate spectrophotometer. The data from two independent experiments done in triplicates was used to calculate the CC50 by nonlinear regression using GraphPad Prism V5.0 software.

### Western blots

The cells were washed with PBS and then lysed using RIPA buffer (1% Triton X-100, 150 mM NaCl, 5 mM EDTA, and 50 mM Tris, pH 7.5) for 30 min at 4°C. The cell lysates were collected after centrifugation at 14,000 × g for 10 min. The proteins were separated on 7% tris-glycine or 8% tricine gels by SDS-PAGE and transferred to a PVDF membrane (Perkin Elmer). The proteins were revealed using a V5-monoclonal antibody (V5-mAb V2660; 1:5000; Invitrogen), ACE2 antibody (rabbit monoclonal ab108252; 1:3,000; Abcam), Actin antibody (rabbit polyclonal A2066; 1:5,000; Sigma), or HA-HRP antibody (12-013-819; 1:3,500; Roche). The antigen-antibody complexes were visualized using appropriate HRP conjugated secondary antibodies and enhanced chemiluminescence kit (ECL; Amersham or Bio-Rad) and normalization was reported to β-actin. Quantification of immune-bands was performed using Image Lab software (Bio-Rad). For analysis of SARS-CoV-2 S virions or pseudovirions, protein extracts of purified viral particles and corresponding producing cells (Calu-3 or 293T17, respectively) were resolved on 10% tris-glycine gels and immunoblotted for spike, nucleocapsid, HIV-1 Gag p24 or actin using anti-V5 (for pseudovirion detection; V2660)/anti-S2 (for virion detection; Sino Biologicals; 40590-T62), anti-N (Sino Biologicals; 40143-MM05), anti-p24 (MBS Hybridoma line 31-90-25) or anti-actin (MP Biomedicals, SKU 08691001), respectively.

### Glycosidase treatment

30 to 50 μg proteins were digested for 90 min at 37°C with endoglycosidase-H (Endo-H; P0702L) or endoglycosidase-F (Endo-F; P0705S) as recommended by the manufacturer (New England Biolabs).

### Inhibitor treatment

At 24h post transfection, cells were incubated for 6h with two pan-PC inhibitors: the cell permeable decanoyl-RVKR-chloromethylketone (cmk; 50 mM; 4026850.001; Bachem) or with the cell surface PC-inhibitor hexa-D-arginine (D6R; 20 μM; 344931; EMD). Culture media were then replaced with fresh ones containing the inhibitors for an additional 24h. For the selective cell-permeable Furin-like inhibitors (BOS; Boston Pharmaceuticals), the cells were treated with the inhibitors at the specified concentration starting at 5h pre-transfection and throughout the duration of the experiment.

### Cell-to-cell fusion assay

HeLa or HeLa TZM-bl cells were plated at 200,000 cells in 12-well plates. HeLa cells were transiently transfected with different constructs of SARS-CoV-2 spike or NL4.3-HIV Env, or an empty vector and 0.2 μg of CMV-Tat plasmid. HeLa TZM-bl cells were transfected with human ACE2, TMPRSS2 or a combination of both. At 6h post-transfection, media were replaced with fresh ones containing Furin-inhibitors, and 24h later the cells were detached with PBS-EDTA (1 µM). Different combinations of HeLa and HeLa-TZM-bl cells were placed in co-culture plate at a ratio of 1:1 for a total of 60,000 cells/well of a 96 well place. After 18-24h the media were removed and 50 μl of cell lysis reagent was added in each well. 20 μl of the cell lysate was used for luciferase reading using 50 μl of Renilla luciferase reagent (Promega, Madison, WI, USA). Relative light units (RLU) were measured using a Promega GLOMAX plate reader (Promega, Madison, WI, USA) and values were reported as fold increase over the RLU measured in co-culture of HeLa cells transfected an empty vector (V) with respective TZM-bl cells.

### Protein immunoprecipitation from co-culture media

When indicated, a secreted form of double tagged spike-glycoprotein of SARS-CoV-2 (N-terminal HA tag and C-terminal V5 tag) from media of co-cultured HeLa and HeLa-TZM-bl cells was analyzed by immunoprecipitation. Namely, 0.3 ml of media were precipitated with 25 µl EZ view Red anti-HA affinity gel (E 6779; Sigma-Aldrich) according to the manufacturers’ protocol. Upon SDS-PAGE separation and PVDF transfer, the proteins were detected using an HA-HRP antibody (12-013-819; 1:3,500; Roche).

### Microscopy

In our luciferase assay, cell co-cultures were plated on glass coverslips. After 18-24h, the cells were incubated with 488 CellMask™ to stain the membrane and then fixed with 4% PFA for 15 min at 4°C. The glass coverslips were mounted on glass slides using ProLong™ Gold Antifade containing DAPI (Invitrogen). The number of syncytia were counted over 10 fields.

### Immunofluorescence

Cell culture and transfection were performed on glass coverslips. Cells were washed twice with PBS and fixed with fresh 4% paraformaldehyde for 10 min at room temperature. Following washes, cells were either non-permeabilized or permeabilized with 0.2% Triton X-100 in PBS containing 2% BSA for 5 min, washed, and then blocking was performed with PBS containing 2% BSA for 1h. Cells were incubated with primary antibodies overnight at 4°C using an antibody against V5 (mouse monoclonal R960-25; 1:1000; Invitrogen), spike (mouse monoclonal GTX632604; 1:500; GeneTex) and ACE2 (goat polyclonal AF933; 1:500; R&D Systems). Following wash, corresponding species-specific Alexa-Fluor (488 or 555)-tagged antibodies (Molecular Probes) were incubated for 1h at room temperature. Coverslips were mounted on a glass slide using ProLong Gold Reagent with DAPI (P36935, Life Technologies). Samples were visualized using a confocal laser-scanning microscope (LSM710, Carl Zeiss) with Plan-Apochromat 63x/1.40 Oil DIC M27 objective on ZEN software.

### Pseudovirus entry

HEK293T-ACE2 or Calu-3 (10,000 cells/well plated in a 96-well dish 24 or 48h before, respectively) were incubated with up to 200 µl filtered pseudovirions for overnight. In certain experiments, target cells were pretreated with BOS-318 (1 µM) for 6h and/or Camostat (40 µM) for 2h before transduction. The overnight incubation with pseudovirions was performed in the presence of the inhibitors. Viral inoculum was removed, then fresh media were added, and the cells cultured for up to 72h. Upon removal of spent media, 293T-ACE2 and Calu-3 cells were gently washed twice with PBS and analyzed for firefly- or nano-luciferase activity, respectively using Promega luciferase assay (Cat # E1501) or Nano-Glo luciferase system (Cat # N1110), respectively.

### Replication competent SARS-CoV-2 Viruses

SARS-CoV-2, which served as the viral source, was originally isolated from a COVID-19 patient in Quebec, Canada and was designated as LSPQ1. The clinical isolate was amplified, tittered in Vero E6 using a plaque assay as detailed below, and the integrity of the S-protein multi-basic protein convertase site validated by sequencing. All experiments involving infectious SARS-CoV-2 virus were performed in the designated areas of the Biosafety level 3 laboratory (IRCM) previously approved for SARS-CoV-2 work.

### Plaque assay in Vero E6 cells

Vero E6 cells (1.2 x 10^5^ cells/well) were seeded in quadruplicate in 24-well tissue culture plates in DMEM supplemented with 10% FBS two days before infection. Cells were infected with up to six ten-fold serial dilutions (10^-2^-10^-6^) of viral supernatant containing SARS-CoV-2 for 1h at 37⁰C (200 µl infection volume). The plates were manually rocked every 15 min during the 1-hour period. Subsequently, virus was removed, cells were washed and overlaying media (containing 0.6% low melt agarose in DMEM with 10% FBS) was added and incubated undisturbed for 60-65h at 37⁰C. Post incubation, cells were fixed with 4% formaldehyde and stained with 0.25% crystal violet (prepared in 30% methanol). High quality plaque pictures were taken using a high resolution DLSR camera (Nikon model: D80, objective: “AF Micro-Nikkor 60mm f/2.8D”). Plaques were counted manually and in parallel, imaged plaque plates were processed and plaques enumerated using an automated algorithm based Matlab software. Virus titer is expressed as plaque-forming units per ml (PFU/ml): (number of plaques x dilution factor of the virus) x 1000 / volume of virus dilution used for infection (in µl). Multiplicity of infection (MOI) expressed as: MOI = PFU of virus used for infection / number of cells.

### Cell infections with fully replicative SARS-CoV-2

Vero E.6 and Calu-3 cells were seeded in duplicates in 12-well plates (2.3 x 10^5^ cells/well) the day before. Cells were pre-treated with various concentrations (0.1-1µM) of BOS-inhibitors and vehicle alone (DMSO) for up to 24h. In certain experiments, Calu-3 were also pre-treated with Camostat for 1h. Thereafter, the cells were infected with SARS-CoV-2 virus at MOI of 0.001 for 1h (Vero E6) or 0.01 for 3h (Calu-3 cells) in 350 µl of serum-free DMEM at 37⁰C with occasional manual rocking of plates. Cells plus media only were used as a control. After incubation, virus was removed, and the cell monolayer was washed twice successively with PBS and serum-free DMEM. New media (total 1ml) containing the concentrations of BOS-inhibitors was subsequently added to cells. Cell-free supernatant (250 µl) was removed at 12, 24 and 48h post infection. The drugs were replenished for 1 ml media at 24h post-infection. The virus supernatants were stored at −80°C until further use. Viral production in the supernatant was quantified using a plaque assay on Vero E6.1 cells as described above. In certain experiments, viral supernatants were harvested at the end of infection and purified on a 20% sucrose cushion using ultracentrifugation as described above. The resulting concentrated virus and corresponding infected cells were analyzed by Western blotting as appropriate.

#### Quantification and statistical analysis

Virus titers quantified by plaque assay in triplicate were shown as mean ± standard deviation. The results from experiments done with two biological replicates and two technical replicates in triplicates were used to calculate the IC_50_ by nonlinear regression using GraphPad Prism V5.0 software. The difference between the control cells (virus with 0.001% DMSO) and the cells treated with BOS-inhibitors were evaluated by Student’s t test. The P values of 0.05 or lower were considered statistically significant (*, p < 0.05; **, p < 0.01; ***, p < 0.001).

## DATA AVAILABILITY

Source data are provided with this paper. The data that support the findings of this study are preserved at repositories of the Montreal Clinical Research Institute (IRCM), Montreal, QC, Canada and available from the corresponding authors upon reasonable request.

## ACKNOWLEDGEMENTS

This work was supported in part by CIHR Foundation grants (NGS: # 148363) and (ÉAC: # 154324), a Canada Research Chairs in Precursor Proteolysis (NGS: # 950-231335), a CIHR CHAMPS Team Grant # HAL 157986 (NGS and ÉAC), Réseau SIDA maladies infectieuses COVID-19 initiative (ÉAC and NGS), the European score project and ANR Reacting COVID-19 (ED and BC). The authors thank the Quebec public health laboratory for providing the infectious isolate LSPQ1 SARS-CoV-2. We thank Paul Bieniasz for the 293T-ACE2 cell line and the pHIV-1NL4-3ΔEnv-NanoLuc construct. The following reagents were obtained from the NIH AIDS Reagent Program, Division of AIDS, NIAID, NIH: TZM-bl cells, from John C. Kappes, Xiaoyun Wu, and Tranzyme, Inc. and HIV-1 pNL4-3 ΔEnv Vpr Luciferase Reporter Vector (pNL4-3.Luc.R-E-) obtained from Nathaniel Landau. We are thankful to Dominic Filion for developing the algorithm for image-assisted plaque quantification. JJ is supported by the CIHR Postdoctoral Fellowship (HIV-435243-73284). We also thank Dr Annik Prat (IRCM) for the design of the summary model shown in Fig. 7. Finally, we would like to thank Mrs. Habiba Oueslati and Brigitte Mary for ther excellent editorial help and organization of the manuscript.

## AUTHOR CONTRIBUTIONS

RE made all the original critical experiments revealing the role of the PCs in spike processing and the effect of their inhibitors. JJ performed all the cell assays with infectious SARS-CoV-2. DSR participated in the biochemical characterizations of TMPRSS2 processing of ACE2 and S1. UA performed all cell-to-cell fusion assays. AE made all the mutants used in the work. RMD generated the HeLa-ACE2 cells and prepared all the cells for *ex vivo* analyses. DNH performed all the immunocytochemical experiments. FD and ML performed experiments related to SARS-CoV-2 pseudovirions. AD and PSO performed all the Furin and TMPRSS2 *in vitro* kinetic cleavage analyses of peptides mimicking the S1/S2 and S2’ sites. CM and KW provided the BOS-inhibitors and their characterization. ED made seminal contributions to the possible role of Furin-like enzymes in the processing of the spike-glycoprotein and actively contributed to the conceptualization and writing of the manuscript. TNQP designed, performed, and analyzed experiments related to viral entry and contributed to the writing of the manuscript. EAC (virology) and NGS (biochemistry and cell biology) conceptualized the research program and provided the intellectual contributions and funding for the whole project. All authors actively contributed to the final version of the manuscript.

## SUPPORTING INFORMATION (SI): TABLE 1; FIGURES 1-7

Table 1: **Sequences of the different peptides mimicking the Cov spike cleavage sites that have been tested in the enzymatic assay.** The arrow indicates the expected cleavage site.

SI-Figure 1: **Importance of Furin in the processing of the Spike-glycoprotein.** (**A**) HeLa cells were first transfected with control non-targeting siRNA (siCTL) or siRNA Furin (siFur) at final concentrations of 20 nM, or mock transfected (N) and 24h later, transfected with empty vector (V) or with that coding for a V5-tagged spike-glycoprotein for an additional 48h. Following lysis, proteins were resolved on SDS-PAGE followed by WB with anti-V5 or anti-Furin antibodies. (**B**) HeLa cells transfected with empty vector (V), V5-tagged wild type spike-protein (WT) or its S1/S2 site mutant (µS1/S2) were treated with Endo-F and Endo-H or mock treated (NT) and analyzed as described in panel A. (**C, D**) HeLa cells transfected with V5-tagged wild type spike-protein (WT) or S1/S2 single mutants (**C**) or S2’ single or double mutants (**D**) in the absence (V) or presence of overexpressed Furin were lysed and analyzed by WB.

SI-Figure 2: **Immunocytochemistry of the co-localization of ACE2 and S-protein or µS1/S2-S in HeLa cells.** Immunofluorescence of S-protein (green), WT (S) or µS1/S2, and ACE2 (red) were revealed using the spike S2-antibody GTX632604 in non-permeabilized (NP) conditions or anti-V5 in permeabilized (P) conditions, and ACE2 antibody AF933. The confocal co-localizations are shown in the merged Figures. Scale bar = 10 µm.

SI-Figure 3: **Furin-like inhibitors strongly reduce SARS-CoV-2 infection in Calu-3 cells.** Calu-3 cells were treated with indicated concentrations of (**A**) BOS-857 and (**B**) BOS-981 and infected with SARS-CoV-2 for 24h. Virus titers in the supernatant were determined by plaque assay on VeroE6 cells (mean plaque forming units [PFU] per ml) ± SD of triplicates, ^∗^p < 0.05; ^∗∗^p < 0.01; ^∗∗∗^p < 0.001). The selectivity index (SI) of (**A**) BOS-857, and (**B**) BOS-981 in Calu-3 cells as shown in top right panel was determined by CC_50_/IC_50_. The left y axis indicates the inhibition of virus titer (percent) relative to that of the untreated control group (red). The right y axis indicates the cell viability (percent) relative to that of the untreated control group (green). Representative plaque images of infected Calu-3 cells treated with indicated doses of BOS-inhibitors are shown in the bottom right panel. Color plaques differentiate the lawn (one color gray per well) from individual plaques (independent colors).

SI-Figure 4: **Furin-like inhibitors modestly reduce virus production in SARS-CoV-2-infected Vero E6 cells in a concentration-dependent manner. (A)** Vero E6 cells treated or not with 1µM BOS-318, BOS-857 or BOS-981 were infected with SARS-CoV-2 for up to 45h. Virus titers in the supernatant obtained at 12, 24 and 48 h post infection were determined by plaque assay on Vero E6. A line graph represents results of the triplicate plaque assay (mean PFU/ml ± SD). **(B, C, and D)** Virus released in the supernatant (48 hr post infection) of infected Vero E6 cells treated with indicated concentrations of (B) BOS-318, (C) BOS-857, or (D) BOS-981 were determined by plaque assay (mean ± SD of triplicates, ∗p < 0.05; ∗∗p < 0.01; ∗∗∗p < 0.001).

SI-Figure 5: **Combination of BOS-981 and Camostat reduces SARS-CoV-2 replication.** Calu-3 cells were treated with BOS-981 and/or Camostat (Camo) and infected with SARS-CoV-2 for 24h. Virus titers in the supernatant were determined by plaque assay on VeroE6 (mean PFU/ml ± SD of duplicates, ^∗^p< 0.05. Representative plaque images of infected Calu-3 cells are shown in the bottom panel. Color plaques differentiate the lawn (one color gray per well) from individual plaques (independent colors).

SI-Figure 6: **Cell-to-cell fusion assay: correlation between syncytia formation and luciferase activity.** (**A**) Cell-to-cell fusion between donor cells (HeLa) and acceptor cells (TZM-bl) was evaluated using confocal microscopy. Hela cells transfected with: (**a**) an empty vector (V), or expressing (**b**) HIV-gp160 and Tat, (**c**) SARS-CoV-2 spike, or (**d**) μS1/S2 were co-cultured with TZM-bl cells for 18h and the number of syncytia was examined using CellMask™ to probe for the plasma membrane and Dapi to stain the nuclei. (**B)** Donor cells were transfected with vectors expressing either no protein (V), Tat, WT-spike (S), Tat and WT-spike (Tat + S) or Tat and HIV-gp160 (Tat + gp160). Acceptor cells were transfected with a vector expressing no protein (V), with ACE2 or directly with Tat as a positive control (hatched bar). After 48h, cells were co-cultured for 18h. Luminescence was normalized to the V value arbitrarily set to 1. Data are presented as mean values ± SD (n=3) and a representative experiment is shown. (**C)** Donor cells were transfected with increasing amount of plasmid expressing WT-spike and acceptor cells were transfected with a vector expressing ACE2. After 48h, cells were co-cultured for 18h, and prepared for luminescence or microscopy. Correlation between the number of syncytia counted by microscopy (n=10 per condition) and the luciferase activity was determined, and the calculated correlation coefficient is R^2^=0.87.

SI-Figure 7: **Secretion of S1.** HeLa cells were transiently co-expressed with double-tagged spike protein (N-terminal HA-tag; C-terminal V5-tag), WT (S) or its mutants, µS1/S2 or µAS1/S2, and ACE2 alone or in combination with TMPRSS2, WT (TMPRSS2) or its S441A active-mutant (µTMPRSS2), at a ratio S:ACE2:TMPRSS2 = 1:0.5:0.5. Immunoblot of the 24h conditioned media was first probed for secreted S1, S1’ and S1_L_ (HA-HRP antibody), stripped and next probed for shed ACE2 (sACE2).

