## Supplemental Table 1 and Figures 1-7 for "SARS-CoV-2 spike-glycoprotein processing at S1/S2 and S2’and shedding of the ACE2 viral receptor: roles of Furin and TMPRSS2 and implications for viral infectivity and cell-to-cell fusion"

### Supporting Information (SI): Table 1; SI Figures 1-7

| | S1/S2 Site | S2' Site |
|---|---|---|
| **SARS-CoV 1 (WT)** | YHTVSLLR↓STSQ | |
| **SARS-CoV 2 (WT)** | TNSPRRAR↓SVAS | SKPSKR↓SFIE |
| **SARS-CoV 2 S1/S2 RRAA** | TNSPRRAASVAS | |
| **SARS-CoV 2 S1/S2 ARAA** | TNSPARAASVAS | |
| **SARS-CoV 2 S1/S2 ARAR** | TNSPARARSVAS | |
| **Sars-CoV 2 S2' RRKR** | | SKRRKRSFIE |

Table 1: **Sequences of the different peptides mimicking the CoV spike cleavage sites that have been tested in the enzymatic assay.** The arrow indicates the expected cleavage site.

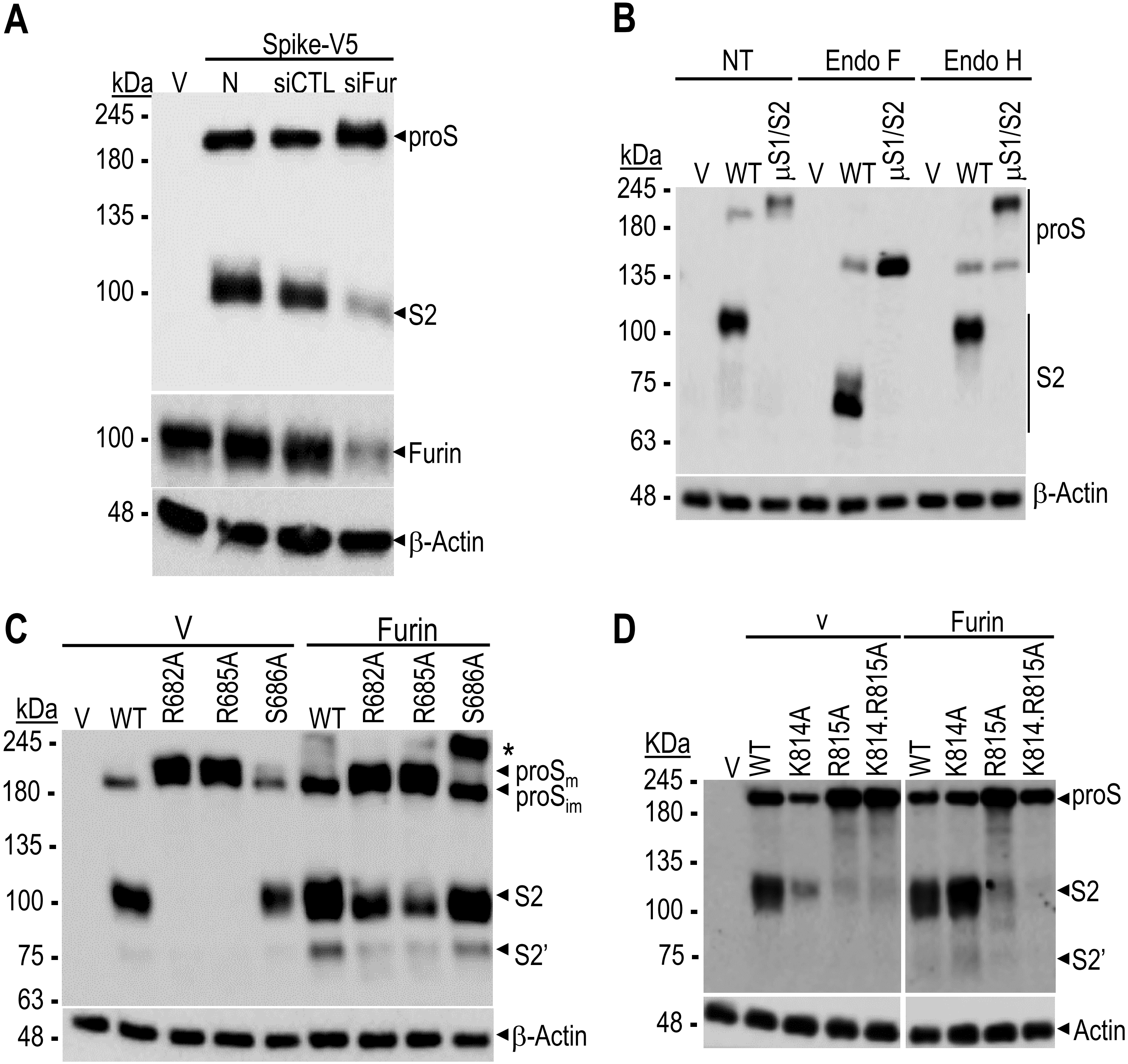

A
Spike-V5
kDa
V
N
siCTL
siFur
245
180
135
100
100
48
proS
S2
Furin
β-Actin
B
NT
Endo F
Endo H
kDa
V
WT
μS1/S2
245
180
135
100
75
63
48
proS
S2
β-Actin
C
V
Furin
kDa
V
WT
R682A
R685A
S686A
WT
R682A
R685A
S686A
245
180
135
100
75
63
48
*
proSm
proSim
S2
S2'
β-Actin
D
v
Furin
KDa
V
WT
K814A
R815A
K814.R815A
WT
K814A
R815A
K814.R815A
245
180
135
100
75
48
proS
S2
S2'
Actin

SI-Figure 1: **Importance of Furin in the processing of the Spike-glycoprotein.** (**A**) HeLa cells were first transfected with control non-targeting siRNA (siCTL) or siRNA Furin (siFur) at final concentrations of 20 nM, or mock transfected (N) and 24h later, transfected with empty vector (V) or with that coding for a V5-tagged spike-glycoprotein for an additional 48h. Following lysis, proteins were resolved on SDS-PAGE followed by WB with anti-V5 or anti-Furin antibodies. (**B**) HeLa cells transfected with empty vector (V), V5-tagged wild type spike-protein (WT) or its S1/S2 site mutant (μS1/S2) were treated with Endo-F and Endo-H or mock treated (NT) and analyzed as described in panel A. (**C, D**) HeLa cells transfected with V5-tagged wild type spike-protein (WT) or S1/S2 single mutants (**C**) or S2' single or double mutants (**D**) in the absence (V) or presence of overexpressed Furin were lysed and analyzed by WB.

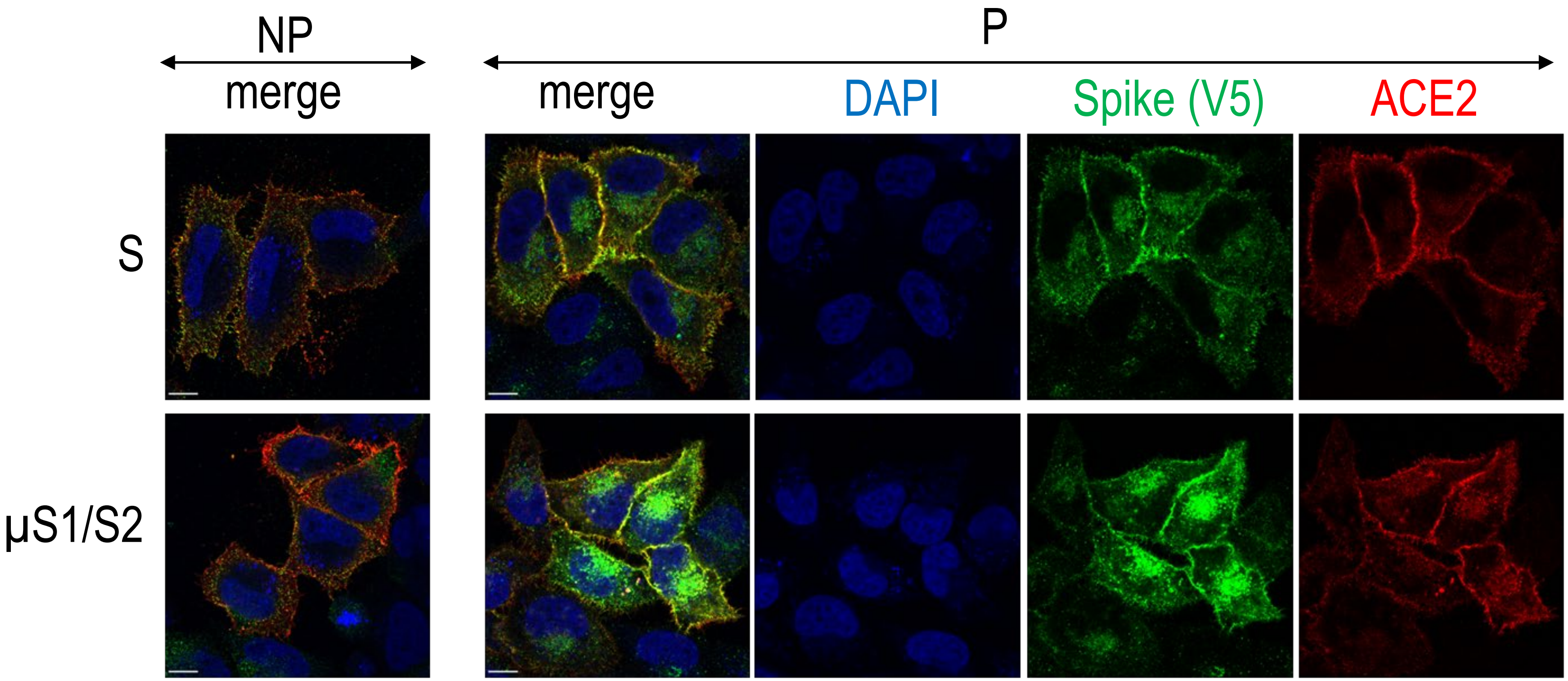


SI-Figure 2: **Immunocytochemistry of the co-localization of ACE2 and S-protein or µS1/S2-S in HeLa cells.** Immunofluorescence of S-protein (green), WT (S) or µS1/S2, and ACE2 (red) were revealed using the spike S2-antibody GTX632604 in non-permeabilized (NP) conditions or anti-V5 in permeabilized (P) conditions, and ACE2 antibody AF933 . The confocal co-localizations are shown in the merged figures. Scale bar = 10 µm.

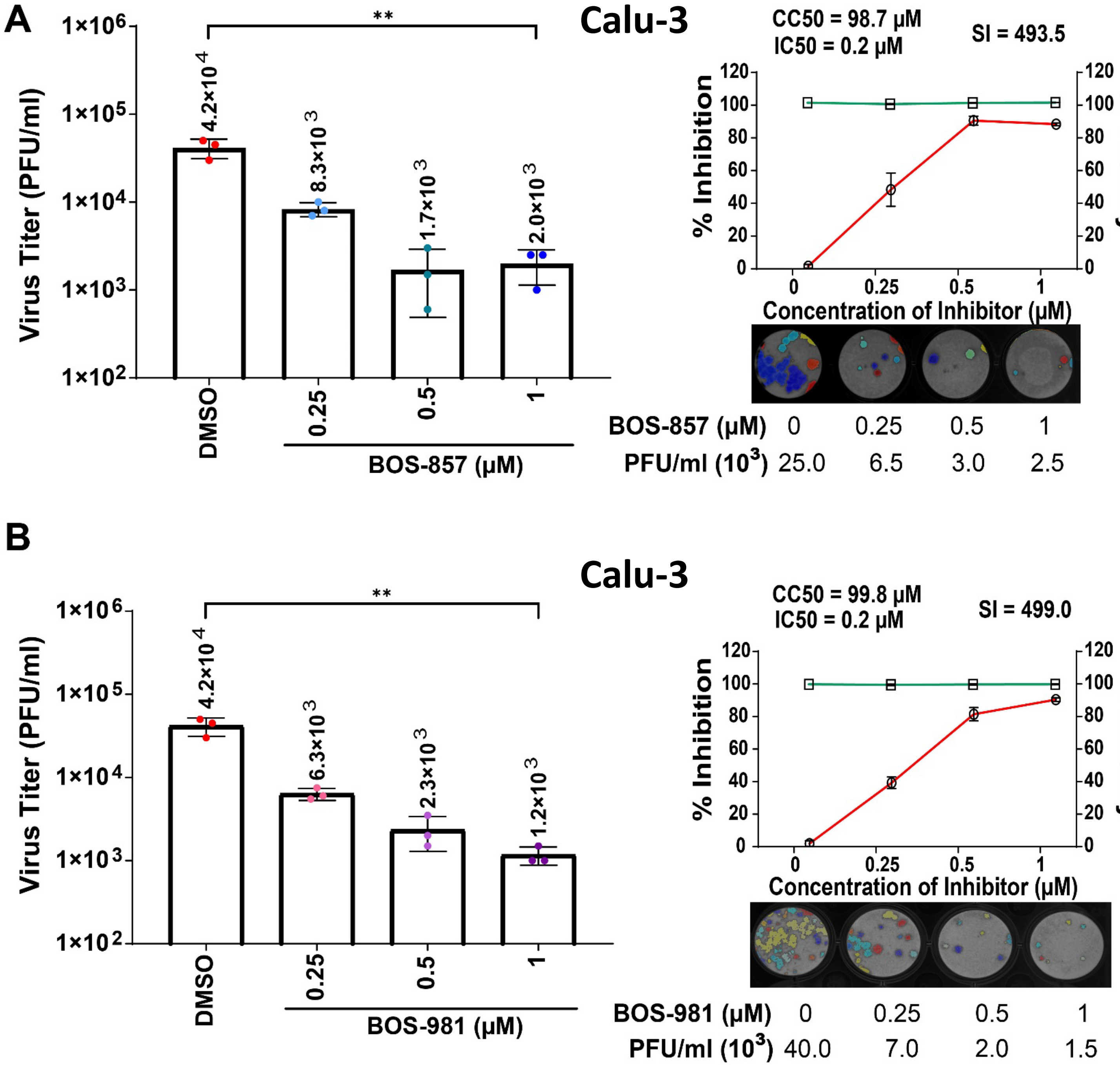
A
Calu-3
Virus Titer (PFU/ml)
1×10^6
1×10^5
1×10^4
1×10^3
1×10^2
**
4.2×10^4
8.3×10^3
1.7×10^3
2.0×10^3
DMSO
0.25
0.5
1
BOS-857 (μM)
CC50 = 98.7 μM
IC50 = 0.2 μM
SI = 493.5
% Inhibition
% Viability
Concentration of Inhibitor (μM)
BOS-857 (μM) 0 0.25 0.5 1
PFU/ml (10^3) 25.0 6.5 3.0 2.5
B
Calu-3
4.2×10^4
6.3×10^3
2.3×10^3
1.2×10^3
BOS-981 (μM)
CC50 = 99.8 μM
IC50 = 0.2 μM
SI = 499.0
BOS-981 (μM) 0 0.25 0.5 1
PFU/ml (10^3) 40.0 7.0 2.0 1.5

SI-Figure 3: **Furin-like inhibitors strongly reduce SARS-CoV-2 infection in Calu-3 cells.** Calu-3 cells were treated with indicated concentrations of (**A**) BOS-857 and (**B**) BOS-981 and infected with SARS-CoV-2 for 24h. Virus titers in the supernatant were determined by plaque assay on VeroE6 cells (mean plaque forming units [PFU] per ml) ± SD of triplicates, $^{*}p < 0.05$; $^{**}p < 0.01$; $^{***}p < 0.001$). The selectivity index (SI) of (**A**) BOS-857, and (**B**) BOS-981 in Calu-3 cells as shown in top right panel was determined by $CC_{50}/IC_{50}$. The left y axis indicates the inhibition of virus titer (percent) relative to that of the untreated control group (red). The right y axis indicates the cell viability (percent) relative to that of the untreated control group (green). Representative plaque images of infected Calu-3 cells treated with indicated doses of BOS-inhibitors are shown in the bottom right panel. Color plaques differentiate the lawn (one color gray per well) from individual plaques (independent colors).

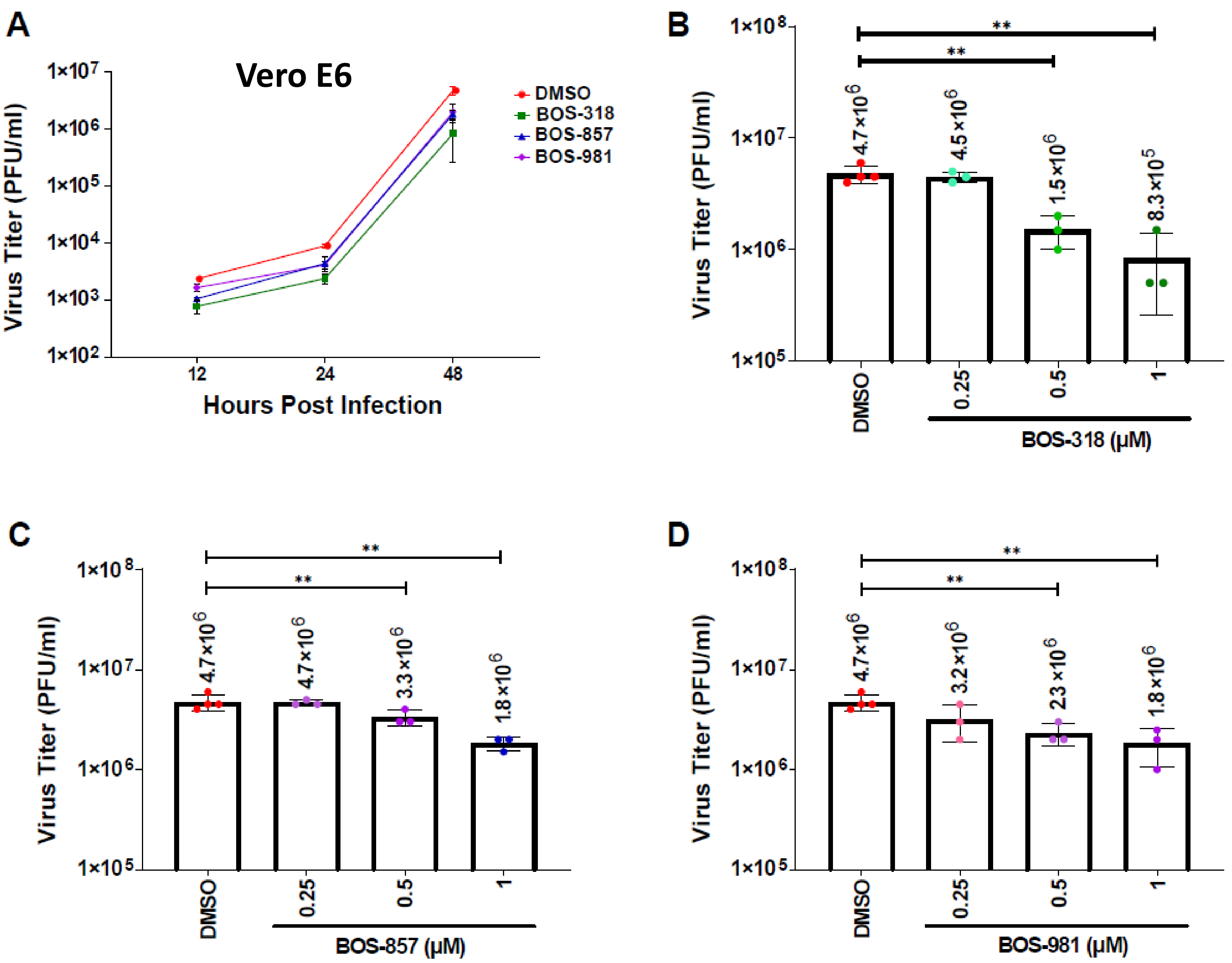
A
Vero E6
Virus Titer (PFU/ml)
Hours Post Infection
DMSO
BOS-318
BOS-857
BOS-981
B
4.7×10^6
4.5×10^6
1.5×10^6
8.3×10^5
BOS-318 (µM)
C
4.7×10^6
4.7×10^6
3.3×10^6
1.8×10^6
BOS-857 (µM)
D
4.7×10^6
3.2×10^6
23×10^6
1.8×10^6
BOS-981 (µM)

SI-Figure 4: **Furin-like inhibitors modestly reduce virus production in SARS-CoV-2-infected Vero E6 cells in a concentration-dependent manner.** **(A)** Vero E6 cells treated or not with 1μM BOS-318, BOS-857 or BOS-981 were infected with SARS-CoV-2 for up to 45h. Virus titers in the supernatant obtained at 12, 24 and 48 h post infection were determined by plaque assay on Vero E6. A line graph represents results of the triplicate plaque assay (mean PFU/ml ± SD). **(B, C, and D)** Virus released in the supernatant (48 hr post infection) of infected Vero E6 cells treated with indicated concentrations of (B) BOS-318, (C) BOS-857, or (D) BOS-981 were determined by plaque assay (mean ± SD of triplicates, $*p < 0.05$; $**p < 0.01$; $***p < 0.001$).

#### Calu-3

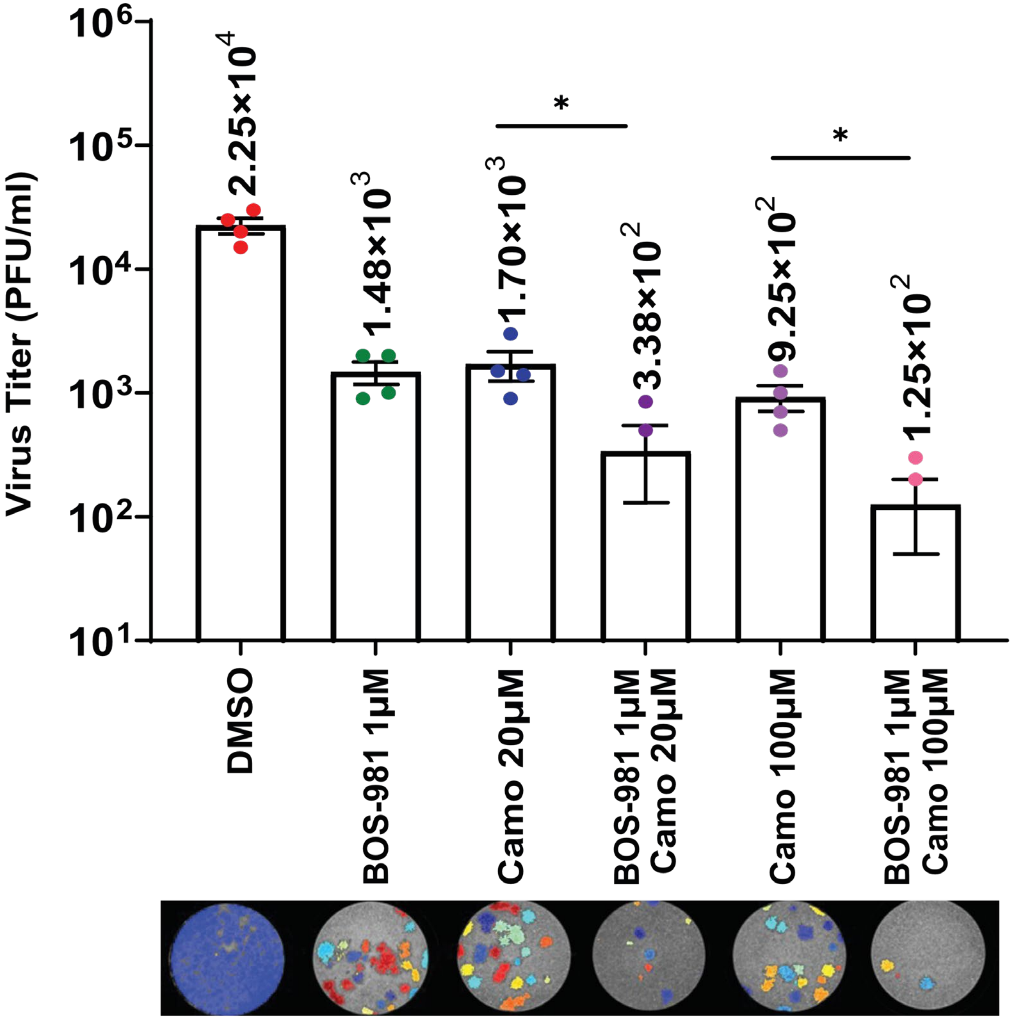


| | | | | | | |
|---|---|---|---|---|---|---|
| BOS-981 (μM) | 0 | 1 | 0 | 1 | 0 | 1 |
| Camostat (μM) | 0 | 0 | 20 | 20 | 100 | 100 |
| PFU/ml ($10^3$) | 25.0 | 1.5 | 1.7 | 0.4 | 1.0 | 0.1 |

SI-Figure 5: **Combination of BOS-981 and Camostat reduces SARS-CoV-2 replication.** Calu-3 cells were treated with BOS-981 and/or Camostat (Camo) and infected with SARS-CoV-2 for 24h. Virus titers in the supernatant were determined by plaque assay on VeroE6 (mean PFU/ml ± SD of duplicates, $^{*}p < 0.05$. Representative plaque images of infected Calu-3 cells are shown in the bottom panel. Color plaques differentiate the lawn (one color gray per well) from individual plaques (independent colors).

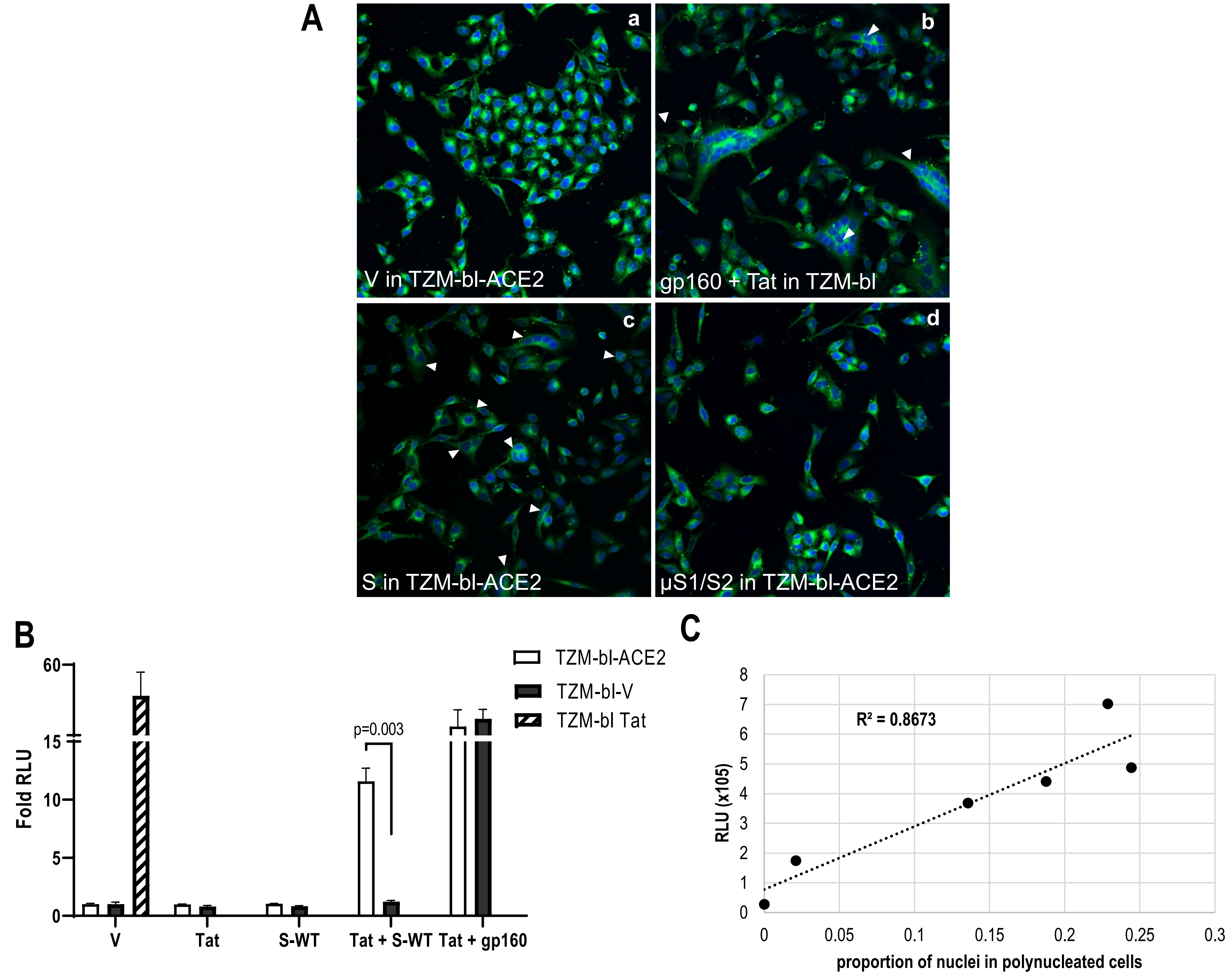

A
a
b
c
d
V in TZM-bl-ACE2
gp160 + Tat in TZM-bl
S in TZM-bl-ACE2
μS1/S2 in TZM-bl-ACE2
B
Fold RLU
60
15
10
5
0
p=0.003
V
Tat
S-WT
Tat + S-WT
Tat + gp160
TZM-bl-ACE2
TZM-bl-V
TZM-bl Tat
C
RLU (x105)
R² = 0.8673
8
7
6
5
4
3
2
1
0
0
0.05
0.1
0.15
0.2
0.25
0.3
proportion of nuclei in polynucleated cells

SI-Figure 6: **Cell-to-cell fusion assay: correlation between syncytia formation and luciferase activity.** (**A**) Cell-to-cell fusion between donor cells (HeLa) and acceptor cells (TZM-bl) was evaluated using confocal microscopy. Hela cells transfected with: (**a**) an empty vector (V), or expressing (**b**) HIV-gp160 and Tat, (**c**) SARS-CoV-2 spike, or (**d**) µS1/S2 were co-cultured with TZM-bl cells for 18h and the number of syncytia was examined using CellMask™ to probe for the plasma membrane and Dapi to stain the nuclei. (**B)** Donor cells were transfected with vectors expressing either no protein (V), Tat, WT-spike (S), Tat and WT-spike (Tat + S) or Tat and HIV-gp160 (Tat + gp160). Acceptor cells were transfected with a vector expressing no protein (V), with ACE2 or directly with Tat as a positive control (hatched bar). After 48h, cells were co-cultured for 18h. Luminescence was normalized to the V value arbitrarily set to 1. Data are presented as mean values ± SD (n=3) and a representative experiment is shown. (**C)** Donor cells were transfected with increasing amount of plasmid expressing WT-spike and acceptor cells were transfected with a vector expressing ACE2. After 48h, cells were co-cultured for 18h, and prepared for luminescence or microscopy. Correlation between the number of syncytia counted by microscopy (n=10 per condition) and the luciferase activity was determined, and the calculated correlation coefficient is $R^2$=0.87.

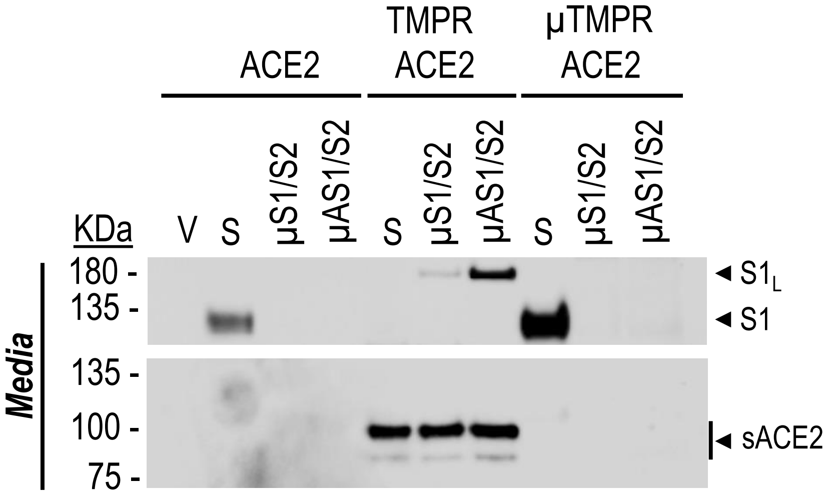


SI-Figure 7: **Secretion of S1.** HeLa cells were transiently co-expressed with double-tagged spike protein (N-terminal HA-tag; C-terminal V5-tag), WT (S) or its mutants, µS1/S2 or µAS1/S2, and ACE2 alone or in combination with TMPRSS2, WT (TMPRSS2) or its S441A active-mutant (µTMPRSS2), at a ratio S:ACE2:TMPRSS2 = 1:0.5:0.5. Immunoblot of the 24h conditioned media was first probed for secreted S1 and S1L (HA-HRP antibody), stripped and next probed for shed ACE2 (sACE2).
